# Dissociable contributions of the amygdala and ventral hippocampus to stress-induced changes in defensive behavior

**DOI:** 10.1101/2023.02.27.530077

**Authors:** Zachary T. Pennington, Alexa R. LaBanca, Patlapa Sompolpong, Shereen D. Abdel-Raheim, Bumjin Ko, Zoe Christenson Wick, Yu Feng, Zhe Dong, Taylor R. Francisco, Madeline E. Bacon, Lingxuan Chen, Sasha L. Fulton, Ian Maze, Tristan Shuman, Denise J. Cai

**Affiliations:** Nash Family Department of Neuroscience, Icahn School of Medicine at Mount Sinai; Department of Pharmacological Sciences, Icahn School of Medicine at Mount Sinai; Howard Hughes Medical Institute, Icahn School of Medicine at Mount Sinai

**Keywords:** Basolateral amygdala, ventral hippocampus, fear, anxiety, PTSD, stress-enhanced fear learning

## Abstract

**Background:** Severe stress can produce multiple persistent changes in defensive behavior relevant to psychiatric illness. While much is known about the circuits supporting stress-induced associative fear, how stress-induced circuit plasticity supports non-associative changes in defensive behavior remains unclear.

**Methods:** Mice were exposed to an acute severe stressor, and subsequently, both associative and non-associative defensive behavioral responses were assessed. A mixture of local protein synthesis inhibition, pan-neuronal chemogenetic inhibition, and projection-specific chemogenetic inhibition were utilized to isolate the roles of the basolateral amygdala (BLA) and ventral hippocampus (vHC) to the induction and expression of associative and non-associative defensive behavioral changes.

**Results:** Stress-induced protein synthesis in the BLA was necessary for enhancements in stress sensitivity but not enhancements in anxiety-related behaviors, whereas protein synthesis in the vHC was necessary for enhancements in anxiety-related behavior but not enhancements in stress sensitivity. Like protein synthesis, neuronal activity of the BLA and vHC were found to differentially support the expression of these same defensive behaviors. Additionally, projection-specific inhibition of BLA-vHC connections failed to alter these behaviors, indicating that these defensive behaviors are regulated by distinct BLA and vHC circuits. Lastly, contributions of the BLA and vHC to stress sensitivity and anxiety-related behavior were independent of their contributions to associative fear.

**Conclusions:** Stress-induced plasticity in the BLA and vHC were found to support dissociable non-associative behavioral changes, with BLA supporting enhancements in stress sensitivity and vHC supporting increased anxiety-related behavior. These findings demonstrate that independent BLA and vHC circuits are critical for stress-induced defensive behaviors, and that differential targeting of BLA and vHC circuits may be needed in disease treatment.

## INTRODUCTION

In immediate response to stressful and life-threatening events, animals display evolutionarily conserved defensive responses, including changes in heart rate and respiration, stress hormone release, as well as the behavioral initiation of fight, flight, and freezing ^1–6^. If sufficiently strong, stressful events can also instantiate persistent changes in how animals interact with their environment. Perhaps most extensively studied are associative fear responses, in which animals engage in defensive behaviors such as freezing when re-exposed to environmental cues present at the time of the initial stressful experience ^1,7–12^. However, after severe stress, animals also display alterations in foraging and exploration in uncertain environments (often referred to as anxiety-related behavior) ^3,5,13,14^, as well as heightened responses to future stressful events ^14–17^. These long-lasting defensive behavioral changes are fundamental to anxiety disorders, which include fear of stress-related cues, heightened stress responses, and reduced environmental engagement; and which are frequently predated by the experience of severe stress ^18–21^.

It is often assumed that many of the defensive behavioral changes observed in the aftermath of stress are fundamentally associative in nature – animals could either be responding to cues that were directly present at the time of stress, or stimuli resembling these cues to some degree (i.e., stimulus generalization) ^22–30^. For example, it is well-documented that startle responses are potentiated by the presence of associative fear cues ^31–34^, suggesting that associative stimuli may drive heightened responses to aversive events after stress. Moreover, it is possible that following stress, alterations in exploration in anxiety-related behavior tests such as the elevated-plus maze could be accounted for by shared features with the environment in which the stressor took place. Lastly, several reports document altered associative fear learning and generalization in humans with anxiety disorders ^23,24,33,35–37^. In light of these findings, broad emphasis has been placed on associative learning processes governing the lasting consequences of stress. However, the explanatory reach of an associative framework has its limits. Pre-weanling rodents incapable of forming associative fear memories nevertheless display increased anxiety-related behavior and heightened responses to subsequent aversive experiences in adulthood ^14^. Moreover, extinguishing fear of stress-associated cues does not necessarily mitigate sensitized responses to new stressors ^15,38,39^. These findings highlight the persistence of some stress-induced behavioral phenotypes despite weak associative fear, indicating a potential dissociation. As such, it could be the case that multiple memory systems – associative and non-associative – support the enduring consequences of stress on defensive behavior. However, a direct biological dissociation of such memory systems has remained elusive. If discovered, this would have broad implications for the treatment of anxiety disorders, potentially explaining why treatments focused on associative processes are ineffective in some individuals ^40–42^.

Here, we explore the contributions of stress-induced plasticity within the ventral hippocampus (vHC) and basolateral amygdala (BLA) to the enduring impacts of stress on associative and non-associative defensive behaviors. Neuronal activity within both the BLA and vHC are well known to regulate defensive behaviors ^43–55^. However, whether stress-induced plasticity within these structures act in concert to support a common defensive behavioral process, or whether they support distinct defensive behavior changes, is unclear. Furthermore, a direct comparison of these structures’ contribution to associative and non-associative defensive processes is lacking. We find that plasticity and neuronal activity within the BLA and vHC support separate non-associative defensive behavior changes in response to stress.

### Methods

Please see the Supplement for detailed experimental procedures.

### Animals

Adult male and female C57BL/6J mice, aged 2-6 months, were used. Experimental procedures were approved Mount Sinai’s IACUC.

### Behavioral testing

#### Trauma and trauma recall

Animals were transported from the vivarium to the experimental testing room and placed in a chamber consisting of unique auditory, olfactory, visual, and spatial cues. During trauma, after a 5 min baseline period, animals received 10, 1 sec, 1 mA, scrambled foot-shocks, with an inter-shock interval of 30 sec. For trauma recall sessions, animals were transported to the same chamber for an 8 min test session.

#### Novel stressor and novel stressor recall

Animals were transported from the vivarium to a chamber with distinct cues from that of the trauma context. After a 3 min baseline period, animals were exposed to a single loud auditory stimulus (3 sec, 130 dB white noise). For novel stressor recall sessions, animals were transported to the same chamber for an 8 min test.

#### Exploratory anxiety-related tests

All anxiety-related behavior tests (light-dark, EPM, and open field) were 5 min in duration.

### Surgery

100-150 nL virus was infused into the BLA (AP: −1.4; ML: 3.3; DV: −5) or vHC (AP: −3; ML: 3.2; DV: −4.5) at 2 nL/sec. For pan-neuronal HM4D experiments, 100 nL of AAV5-hSyn-HM4Di-mCherry (Addgene 50475) or AAV5-hSyn-eGFP (Addgene 50465) was infused. For projection-specific HM4D experiments, 150 nL containing a cocktail of AAVrg-ef1a-Cre (Addgene 55636) and AAV8-hSyn-eGFP (Addgene 50465) were infused into the projection target structure. Additionally, 150 nL of AAV5-hSyn-DIO-HM4Di-mCherry (Addgene 44362) or AAV5-hSyn-DIO-mCherry (Addgene 50459) was infused into the projection origin structure. For cannulation surgeries, 26 gauge guide cannula (P1 Technologies; 8IC315GMNSPC) were implanted overlying the BLA or vHC.

### Drugs

Anisomycin (Sigma A9789) was administered systemically at a dose of 150 mg/kg (10 mL/kg, s.c.) ^101,102^. Because numerous waves of protein synthesis can support memory consolidation ^68,103,104^, we administered anisomycin 3 times, once every 4 hours. Control animals received vehicle at the same times. Anisomycin (10 ng/nL) was infused intracranially at a rate of 150 nL/min. 300 nL of anisomycin/vehicle was administered per hemisphere in the vHC and 200 nL was administered per hemisphere in the BLA. CNO-dihydrochloride (3 mg/kg, i.p., Tocris), was injected 30-40 minutes prior to behavior.

## RESULTS

### Acute severe stress produces multiple lasting changes in defensive behavior

We first established a behavioral protocol in which a single acute stressor produces lasting changes in multiple defensive behaviors, adapting a prior model that has been used in mice and rats ^14,15,56–58^ (Fig 1A). Animals were placed in a distinct environment where they received 10 foot-shocks during a 10-min period (Fig 1B. ‘trauma’, T), or were placed in the same environment but did not receive foot-shocks (‘no-trauma’, NT). A week later, multiple defensive behaviors were assessed. To examine associative fear, animals were returned to the trauma environment (trauma recall). As expected, trauma-exposed animals spent a large amount of time freezing (Fig 1C). In the light-dark box, an exploratory test that captures rodents’ natural avoidance of well-lit places and is sensitive to anxiolytics ^59,60^, trauma-exposed animals showed increased anxiety-related behavior, reflected in more time spent in the dark side of the light-dark box (Fig 1D). Lastly, we assessed the animals’ stress sensitivity by placing the animals in a novel environment, in which they showed very little initial freezing (Fig 1E, left panel), and presenting them with a loud auditory startle stimulus. When returned to this environment the next day, trauma-exposed animals showed substantially more freezing, evidence of stress sensitization (Fig 1E, right panel). Importantly, we demonstrate that all of these defensive behavioral changes – in associative fear, anxiety-related behavior, and stress sensitization – are proportional to the magnitude of the initial trauma (Fig S1). Additionally, although sex-differences are common amongst anxiety disorders ^61,62^, we found no behavioral differences in male and female mice on the dependent variables examined (Fig S1). Lastly, although stress sensitization is often termed stress-enhanced fear learning ^15^, learning curve analyses revealed that enhanced learning likely reflects heightened sensitivity to aversive stimuli, as opposed to an enhanced learning rate (Fig S2).

**Figure 1:**
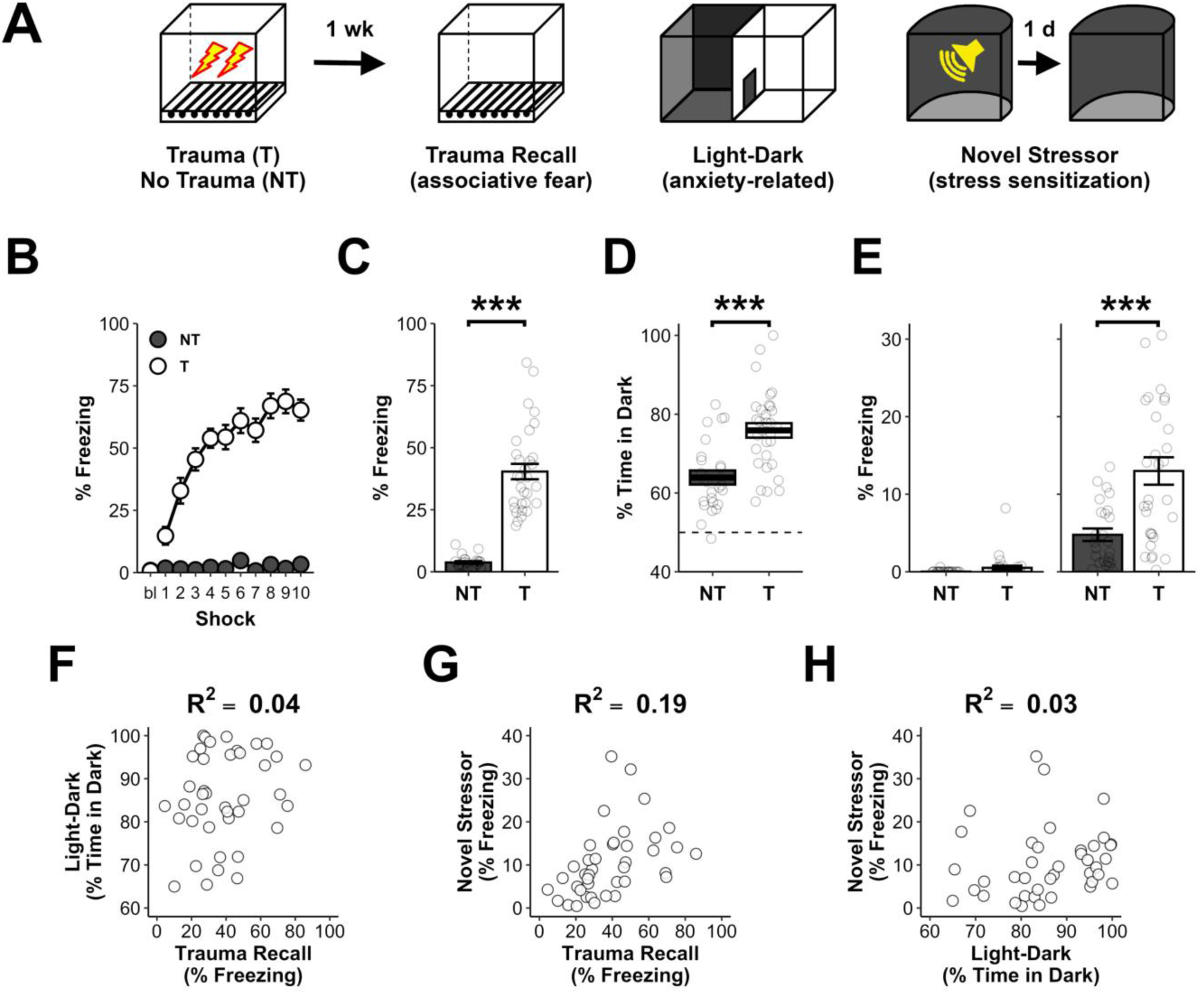
Acute stress produces multiple, lasting, changes in defensive behavior. **A)** Animals were exposed to an environment in which they received 10 foot-shocks (trauma, T) or were placed in the same environment and received no foot-shocks (no-trauma, NT). A week later, they were tested for associative fear of the trauma environment, anxiety-related behavior in the light-dark test, and their response to a novel stressor in a new environment to assess stress sensitization. **B)** Trauma-exposed animals displayed high levels of post-shock freezing during the trauma (Trauma: F_1,33_=167.4, p<0.001). **C)** Trauma-exposed animals displayed strong associative fear of the trauma environment (Trauma: F_1,52_=121.6, p<0.001). **D)** Trauma-exposed animals displayed increased anxiety-related behavior in the light-dark test (Trauma: F_1,52_=19.7, p<0.001). **E)** Trauma-exposed animals did not differ in baseline levels of freezing when initially placed in the environment of the novel stressor (left. Trauma: F_1,52_=2.6, p=0.11), but displayed increased fear of the novel stressor environment when returned to this environment the next day, evidence of stress sensitization (right. Trauma: F_1,52_=16.1, p<0.001). **F)** Correlation between trauma recall and anxiety-related behavior in light-dark test (R^2^=0.04, p=0.19). **G)** Correlation between trauma recall and novel stressor response (R^2^=0.19, p<0.01). **H)** Correlation between anxiety-related behavior in light-dark test and novel stressor response (R^2^=0.13, p=0.32). For B-E, NT=25(13 female) and T=31(16 female) mice. For F-H, T=40 mice. p<.05 (*), p<0.01 (**), p<0.001 (***). Error bars reflect standard error of the mean.

Next, as a preliminary means of addressing if these defensive behaviors convey information about unique biobehavioral processes, we correlated these phenotypes in a large group of trauma-exposed animals (vehicle-treated control animals in Fig 2F-H), as high inter-phenotype correlations would suggest shared biological origins. We found relatively small correlations between behavioral tests, with large amounts of variance in each being unexplained by the others (Fig 1F-H). Though each of these measures has imperfect test-retest reliability, these findings nevertheless suggest that these phenotypes may be independently regulated.

**Figure 2:**
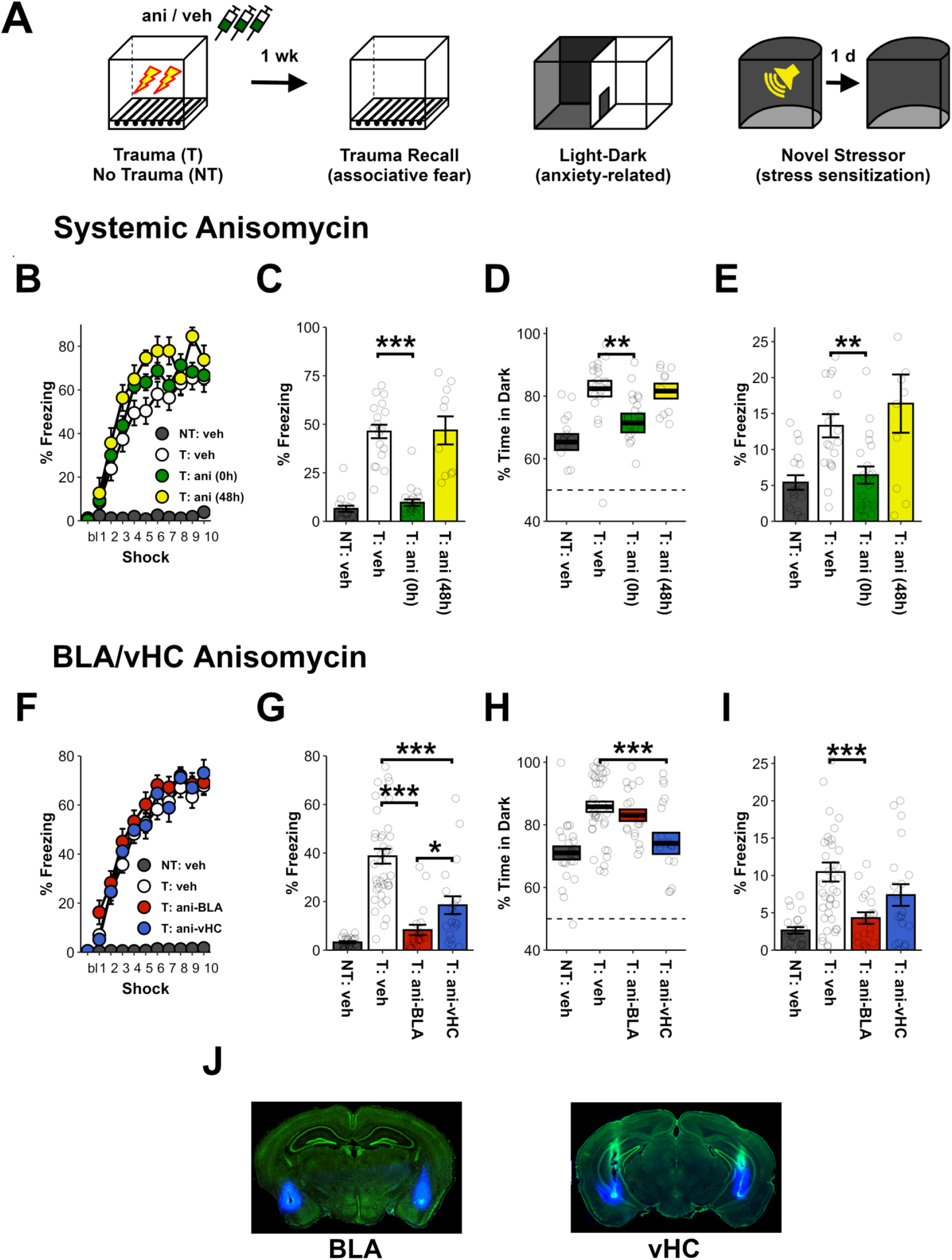
Stress-induced protein synthesis in the BLA and vHC support distinct changes in non-associative defensive behavior. **A)** After trauma (T) or no-trauma (NT), animals were administered 3 injections of anisomycin (ani) or vehicle (veh). A week later, they were tested for associative fear of the trauma environment, anxiety-related behavior in the light-dark test, and their response to a novel stressor in a new environment. For B-E, anisomycin/vehicle was administered systemically either immediately (0h) or 48 hours (48h) after trauma. For F-J, anisomycin/vehicle was administered directly into either the BLA or vHC immediately after trauma. **B)** No differences were observed between trauma-exposed animals during the initial trauma (Group: F_2,43_=2.8, p=0.07; Group x Shock: F_20,430_=0.7, p=0.78). **C)** Anisomycin given systemically immediately after trauma, but not 48 later, reduced associative fear of the trauma environment (NT: veh vs T: veh, t_25_=10.5, p<0.001; T: veh vs T: ani (0h), t_26.1_=9.5, p<0.001; T: veh vs T: ani (48h), t_13.3_=0.1, p=0.95). **D)** Anisomycin given systemically immediately after trauma, but not 48 later, reduced anxiety-related behavior in the light-dark test (NT: veh vs T: veh, t_33.8_=4.8, p<0.001. T: veh vs T: ani (0h), t_36.1_=2.8, p<0.01 T: veh vs T: ani (48h), t_24.7_=0.2, p=0.83). **E)** Anisomycin given systemically immediately after trauma, but not 48 later, reduced stress sensitization (NT: veh vs T: veh, t_29.7_=4.1, p<0.001; T: veh vs T: ani (0h), t_33.6_=3.4, p<0.01; T: veh vs T: ani (48h), t_12_=0.7, p=0.49). **F)** No differences were observed between trauma-exposed animals during the initial trauma (Group: F_2,76_=1.1, p=0.33; Group x Shock: F_20,760_=0.5, p=0.96). **G)** Anisomycin in the BLA and vHC reduced associative fear of the trauma environment (NT: veh vs T: veh, t_40.7_=11.5, p<0.001; T: veh vs T: ani-BLA, t_56.9_=8.2, p<0.001; T: veh vs T: ani-vHC, t_44_=4.2, p<0.001; T: ani-BLA vs T: ani-vHC: t_30.5_=2.4, p=0.02). **H)** Anisomycin in the vHC, but not the BLA, reduced anxiety-related behavior in the light-dark test (NT: veh vs T: veh, t_47.5_=5.5, p<0.001; T: veh vs T: ani-BLA, t_44.9_=1.1, p=0.28; T: veh vs T: ani-vHC, t_28.2_=3.1, p<0.01). **I)** Anisomycin in the BLA, but not the vHC, reduced stress sensitization relative to controls (NT: veh vs T: veh, t_47.6_=5.8, p<0.001; T: veh vs T: ani-BLA, t_56.4_=4.1, p<0.001; T: veh vs T: ani-vHC, t_46.2_=1.6, p=0.12). **J)** Example placement of cannula injectors in the BLA and vHC for intracranial infusions. For systemic injections (B-E), NT: veh=17(5 female), T:veh=19(5 female), T:ani (0h)=20(5 female), and T:ani (0h)=10(5 female) mice. For intracranial infusions (F-J), NT:veh=23, T:veh=40, T:ani-BLA=19, and T:ani-vHC=20 mice. Half of vehicle-treated animals had cannula in BLA and the other half in the vHC. P<.05 (*), p<0.01 (**), p<0.001 (***). Error bars reflect standard error of the mean. Statistics in main text.

### Stress-induced protein synthesis in the BLA and vHC produce distinct changes in non-associative defensive behavior

In order to assess how stress-induced plasticity supports persistent changes in defensive behavior, we utilized post-stress administration of the protein synthesis inhibitor anisomycin, as protein synthesis is known to support the consolidation of many forms of memory and synaptic plasticity ^63–69^. Furthermore, because manipulations of protein synthesis can be done after a learning experience, they provide a means of disrupting the consolidation of an experience without altering its initial encoding, and also shed light on a translationally relevant time period for trauma intervention.

To validate that the emergence of the observed defensive behavioral changes are indeed supported by stress-induced protein synthesis, we first assessed the effects of systemically administering anisomycin at various time points after trauma (Fig 2B-E). Animals underwent trauma and were given anisomycin either immediately after trauma (T: ani (0h)), 48 hours after trauma (T: ani (48h)), or underwent trauma but were injected with vehicle (T: veh). Control mice were placed in the same environment but did not receive trauma and were treated with vehicle (NT: veh). During the initial trauma, groups receiving trauma did not differ in their level of freezing, indicating that there were no pre-existing group differences (Fig 2B). A week later, trauma-exposed animals treated with vehicle exhibited associative fear in the trauma recall test (Fig 2C), increases in anxiety-related behavior in the light dark-test (Fig 2D), and heightened fear of the novel stressor environment (Fig 2E), relative to no-trauma controls. Anisomycin administration immediately after trauma reduced all of these stress-induced defensive behaviors relative to trauma-exposed animals given vehicle (Fig 2C-E). However, anisomycin given 48 hours after trauma did not reduce defensive behaviors relative to trauma-exposed animals given vehicle (Fig 2C-E). Therefore, protein synthesis occurring just after trauma appears critical to the induction of the observed defensive phenotypes.

Next, we assessed the impacts of targeting trauma-induced protein synthesis specifically in the BLA or vHC, regions previously linked to regulating defensive behaviors and anxiety disorders (Fig 2F-J) ^70–73^. Mice had cannulas implanted above either the BLA or vHC (Fig 2J. See Fig S3 for placement in all animals). Following surgical recovery, animals then underwent trauma and immediately after received intracranial infusions of anisomycin or vehicle. Alternatively, they experienced no trauma and were treated with vehicle. Animals treated with vehicle in the BLA and vHC showed no behavioral differences and are collapsed here (Fig S4). Prior to vehicle/anisomycin treatment, no differences were observed in freezing during the trauma session for animals that underwent trauma (Fig 2F). Additionally, as anticipated, trauma-exposed animals treated with vehicle exhibited a strong associative memory (Fig 2G), heightened anxiety-related behavior in the light-dark test (Fig 2H), and heightened fear of the novel stressor (Fig 2I), relative to no-trauma controls. These behaviors were differentially affected by blocking trauma-induced protein synthesis in the BLA and vHC. In the trauma recall test, anisomycin in either the BLA or vHC were effective at reducing associative freezing relative to trauma controls, though the BLA appeared to contribute to a more sizable degree (Fig 2G). In the light-dark test, anisomycin in the vHC greatly attenuated trauma-induced increases in anxiety-related behavior (Fig 2H). However, anisomycin in the BLA was without effect (Fig 2H). Lastly, anisomycin in the BLA was able to block the enhanced sensitivity to a novel stressor, whereas anisomycin in the vHC was without effect (Fig 2I). Notably, the dose of anisomycin used here was found to alter memory consolidation and protein synthesis, but not memory expression, suggesting it did not acutely influence neuronal function (Fig S5). Moreover, although the dosing regimen used was found to alter memory consolidation when given immediately after a learning event, it had no long-term deleterious impacts on future learning, indicating that no permanent damage was produced by anisomycin administration (Fig S6).

These findings highlight that while stress-induced protein synthesis in the BLA is paramount for associative fear and heightened stress sensitivity, it is not necessary for alterations in anxiety-related behavior. Conversely, stress-induced protein synthesis in the vHC is essential for increased anxiety-related behavior, and to a lesser extent associative fear recall, but not heightened stress sensitivity. Importantly, the finding that blockade of protein synthesis in the BLA had a profound impairment on associative fear for the trauma environment but no detectable effect on the light-dark test further suggests a dissociation between anxiety-related behavior and associative fear. Similarly, the fact that blockade of protein synthesis in the vHC reduced associative fear but did not alter stress sensitization indicate that these phenotypes are also dissociable.

### Neuronal activity in the BLA and vHC support distinct stress-induced defensive behaviors

The prior findings indicate that stress-induced protein synthesis within the BLA and vHC support the induction of distinct post-stress defensive phenotypes. However, neuronal activity in both regions could potentially still be necessary at a later time-point to express changes in stress sensitivity and anxiety-related behavior. To evaluate this possibility, we next used a chemogenetic system ^74^ to inhibit the BLA or vHC during testing of associative memory recall, anxiety-related behavior, and stress sensitization.

We first verified that administration of the agonist CNO-dihydrochloride (cno) was able to inhibit neuronal activity of cells expressing the HM4D receptor in the BLA/vHC. A pan-neuronal virus expressing HM4D was infused into either the BLA or vHC (Fig 3A). Recording from these neurons in slice ~1 month later, we found robust inhibition of HM4D-expressing cells when cno was applied to the bath (Fig 3B).

**Figure 3.**
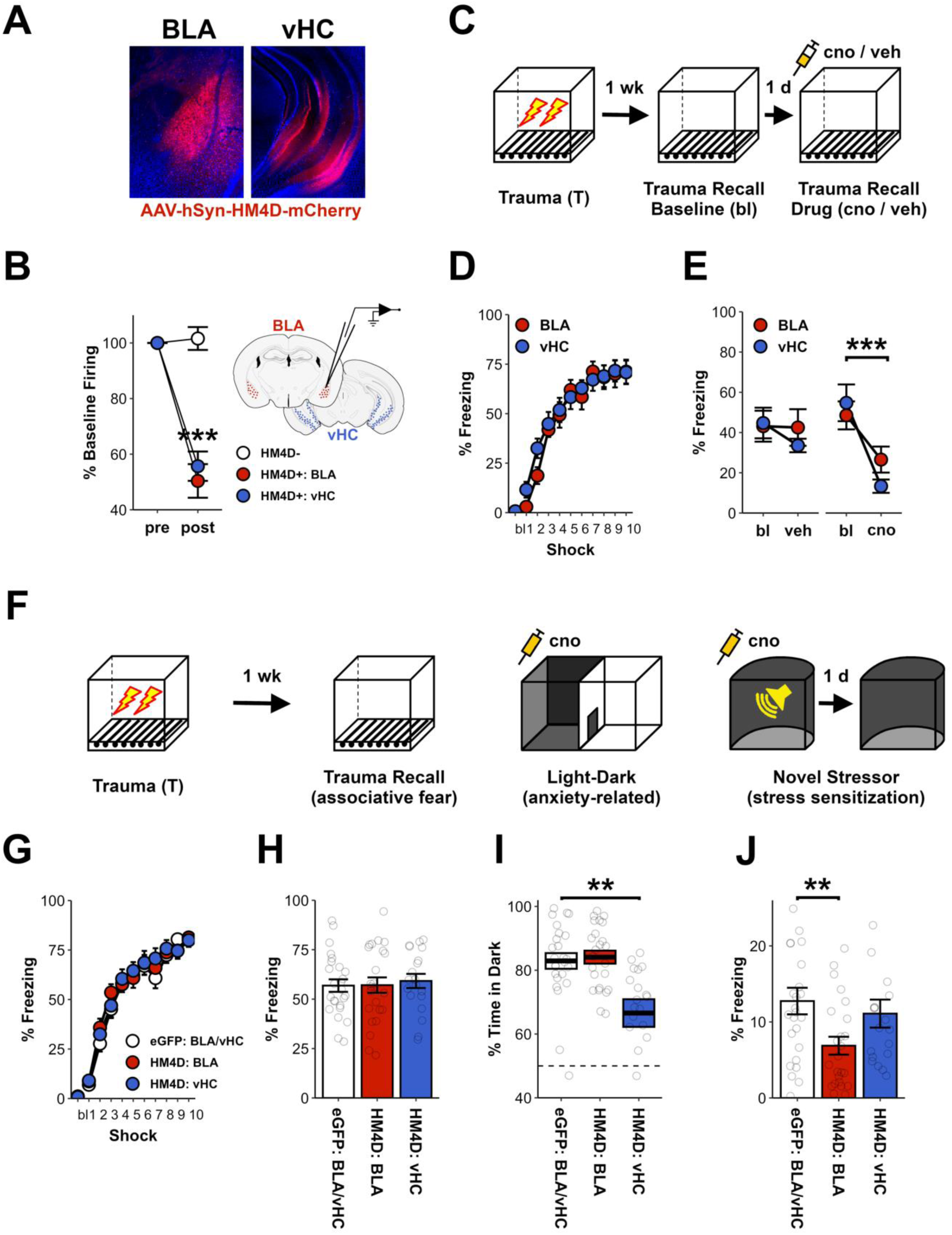
Neuronal activity in the BLA and vHC support distinct stress-induced defensive behaviors. **A)** A pan-neuronal virus expressing the inhibitory chemogenetic receptor HM4D, or eGFP, was infused in the BLA or vHC. **B)** HM4D+ neurons, as well as neighboring HM4D-neurons, were recorded before and after cno application. Application of cno dramatically reduced action potentials in HM4D+ neurons (HM4D+ cells – Pre-Post: F_1,10_=137, p<0.001; Pre-Post X Region: F_1,10_=0.4, p=0.52; HM4D-cells – Pre-Post: F_1,1_= 0.2, p=0.76) **C)** Animals underwent trauma and a week later were tested twice for their associative recall of the traumatic event, first in a drug-free baseline test (bl), and second, after receiving an injection of cno or saline (veh). **D)** Animals with HM4D in the BLA and vHC did not differ during the initial trauma (Region: F_1,28_=0.3, p=0.59; Region x Shock: F_10,280_=0.7, p=0.66). **E)** Administration of cno reduced freezing in animals with HM4D in either the BLA or vHC (cno: F_1,14_=33.5, p<0.001; cno X Region: F_1,14_=3.2, p=0.1; veh: F_1,12_=2.8, p=0.12; veh X Region: F_1,14_=2.2, p=0.16). **F)** Animals underwent trauma and a week later, they were tested for associative fear of the trauma environment, anxiety-related behavior in the light-dark test, and their response to a novel stressor in a new environment. The BLA/vHC were inhibited via cno administration prior to the light dark test, as well as prior to the novel stressor. **G)** No group differences were observed during the initial trauma (Group: F_2,65_=0.2, p=0.82; Group X Shock: F_20,650_=0.6, p=0.84). **H)** No group differences were observed during the drug-free trauma recall test (Group: F_2,65_=0.1, p=0.88). **I)** Inhibition of the vHC, but not the BLA, reduced anxiety-related behavior in the light-dark test (eGFP vs HM4D: BLA, t_45.8_=0.4, p=0.71; eGFP vs HM4D: vHC, t_29.2_=3.3, p<0.01). **J)** Inhibition of the BLA, but not the vHC, reduced freezing in the test of stress sensitization (eGFP vs HM4D: BLA, t_41.7_=2.8, p<0.01; eGFP vs HM4D: vHC, t_40.4_=0.7, p=0.52). For electrophysiological recordings in B, HM4D-=2, HM4D+: BLA=6, and HM4D+: vHC=7 cells. For effects of inhibition on recall in C-E, BLA: veh=7, BLA: cno=7, vHC: veh=7, and vHC: cno=9 mice. For effects of inhibition on light-dark and novel stressor in F-J, eGFP: BLA/vHC=25, HM4D: BLA=24, and HM4D: vHC=19 mice. P<.05 (*), p<0.01 (**), p<0.001 (***). Error bars reflect standard error of the mean. Statistics in main text.

We next tested whether neuronal activity within the BLA and vHC are necessary for trauma memory recall. Animals expressing HM4D in either the BLA or vHC underwent trauma (Fig 3D) and a week later, their associative recall of the event was assessed – first during a drug-free, baseline test, and then, after receiving an injection of saline (veh) or cno (3 mg/kg, i.p.). As expected, inhibition of either BLA or vHC reduced freezing levels relative to baseline (Fig 3E); vehicle-treated animals did not show altered freezing levels (Fig 3E). These findings mirror our protein synthesis results, and are consistent with prior reports that the activity of both the BLA and vHC are important for associative fear^75^.

Then, to assess the contributions of BLA and vHC neural activity to the expression of enhanced anxiety-related behavior and stress sensitivity after trauma, a virus expressing HM4D was infused into the BLA/vHC, or a control virus expressing eGFP was infused. Controls with eGFP in the BLA or vHC were collapsed into a common control group (eGFP: BLA/vHC. See Supplementary Table 1 for comparisons of BLA/vHC controls). A month later, all animals underwent the trauma procedure (Fig 3F). Notably, no behavioral differences were observed during the initial trauma, suggesting that expression of the receptor alone had no effect on the acquisition or expression of conditioned freezing (Fig 3G). Additionally, when placed drug-free in the trauma environment a week later, no group differences in freezing were observed (Fig 3H). Consistent with our finding that stress-induced protein synthesis in the vHC supports enhancements in anxiety-related behavior, inhibition of the vHC produced a dramatic decrease in time spent on the dark side in the light-dark test (Fig 3I), whereas inhibition of the BLA was without effect (Fig 3I). Similarly, in the test of stress sensitization, animals in which the BLA was inhibited froze less than controls (Fig 3J), whereas inhibition of the vHC was without effect (Fig 3J).

To confirm the reliability of these findings, utilizing a different chemogenetic receptor ^76^ we replicated the finding that BLA activity supports heightened stress sensitivity, whereas the vHC supports anxiety-related behavior, and both of these structures support associative memory recall (Fig S8). Moreover, to test whether our finding that the vHC selectively contributes to anxiety-related behavior are generalizable, we tested the impacts of inhibiting the BLA and vHC across anxiety-related behavior tests (light-dark, elevated plus maze, and open field), as each test is likely to tap into slightly different cognitive/behavioral processes. Although slight differences were observed across tests and chemogenetic systems, we broadly found that inhibiting the vHC, but not the BLA, was able to reduce anxiety-related behavior (Fig S9).

In summary, inactivation of the BLA/vHC mirrored the effects observed with protein synthesis inhibition, such that the BLA supports heightened stress sensitivity, and the vHC supports enhanced anxiety-related behavior. Therefore, it appears at both the level of protein synthesis and neuronal activity that stress-induced changes in stress sensitivity and anxiety-related behavior are supported by different neural substrates.

### Inhibiting reciprocal BLA-vHC connections fails to alter stress sensitivity and anxiety-related behavior

Above, we have demonstrated that stress-induced protein synthesis and subsequent neuronal activity within the BLA and vHC support distinct defensive behavioral changes. This is incredibly surprising given that the BLA and vHC share reciprocal monosynaptic connections ^77–79^, and prior reports indicating these projections support at least some defensive behaviors in stress-naïve animals ^79–81^. That said, both the BLA and vHC contain output neurons that project to distinct downstream structures ^77,78,80–82^, and there is evidence that these projections can contribute to different defensive processes ^80^. Therefore, it may be that stress-induced changes in anxiety-related behavior and stress sensitivity are not dependent upon BLA-vHC connectivity. To directly test this possibility, we used a retrograde viral approach to selectively inhibit cells in the BLA that project to the vHC, or vice versa. A cre-expressing retrograde AAV was injected into either the BLA or vHC, and a cre-dependent HM4D virus or a control virus expressing only mCherry was injected into the other structure (Fig 4A. Animals with mCherry in the BLA and vHC were collapsed into a single group. See Supplementary Table 1 for comparisons of BLA/vHC controls). A month later, animals underwent the trauma protocol previously described, and BLA-vHC projections were inhibited during the light-dark test as well as during the novel stressor (Fig 4B). As expected, with both structures ‘online’, groups did not differ in their response to the initial trauma (Fig 4C), nor did they differ in the trauma recall test (Fig 4D). Strikingly, selective inhibition of either projection did not alter anxiety-related behavior in the light-dark test (Fig 4E). Similarly, inhibiting either projection did not alter the response to the novel stressor in the test of stress sensitization (Fig 4E). Consequently, it appears that the reciprocal connections between the BLA and vHC do not play a pivotal role in these specific stress-induced changes in defensive behavior.

**Figure 4.**
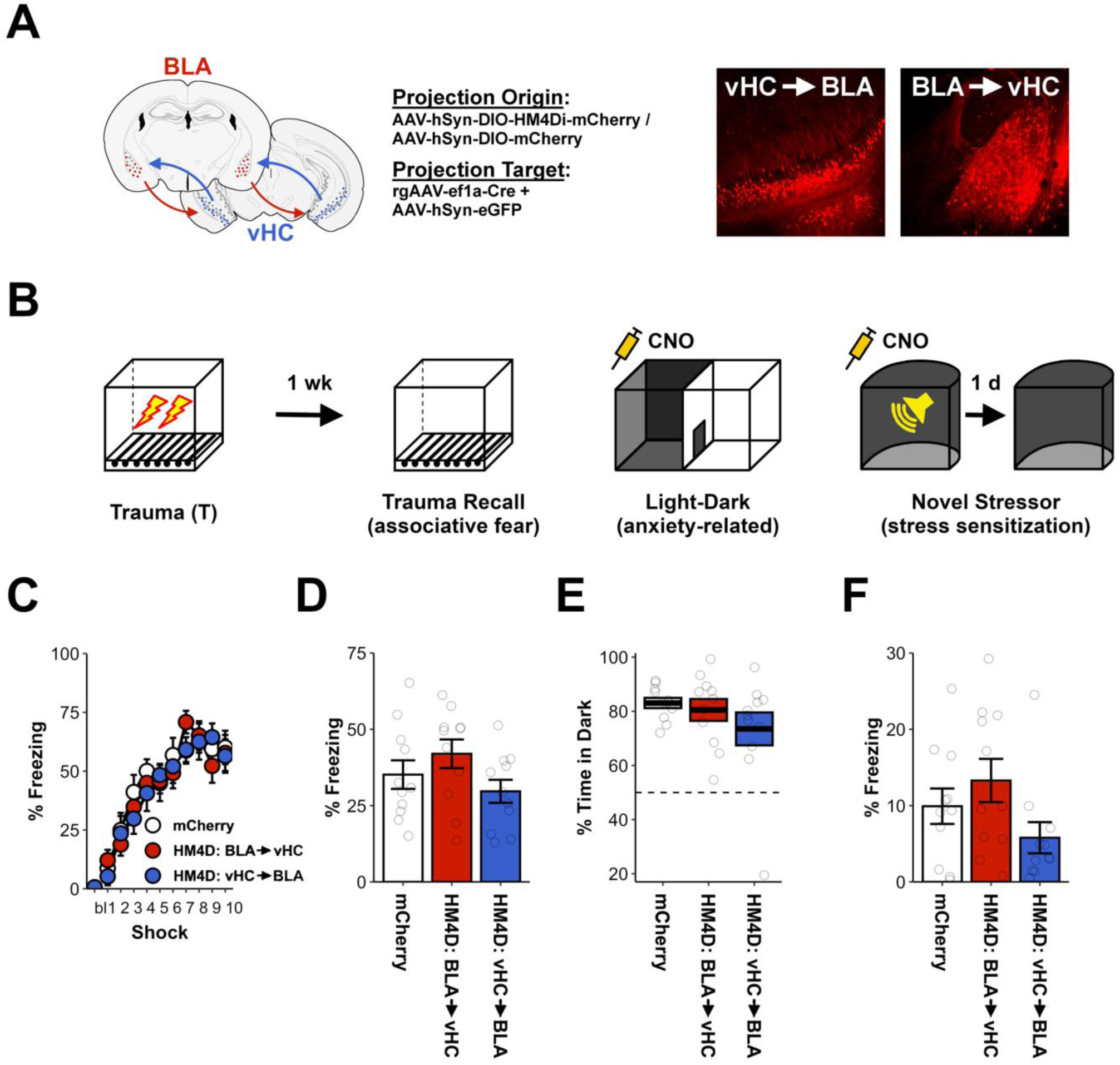
Reciprocal BLA-vHC connections do not support stress-induced changes in anxiety-related behavior or stress sensitivity. **A)** Projection-specific targeting of BLA cells projecting to the vHC, or vice versa, was accomplished by infusing a cre-expressing virus into the projection target structure and cre-dependent HM4D/control virus into the projection origin structure. eGFP-expressing virus was co-infused into the projection target to confirm surgical placement. **B)** Animals underwent trauma and a week later were tested for associative fear of the trauma environment, anxiety-related behavior in the light-dark test, and their response to a novel stressor in a new environment. The BLA-vHC connections were inhibited via cno administration prior to the light dark test, as well as prior to the novel stressor. **C)** No group differences were observed during the initial trauma (Group: F_2,30_=0.1, p=0.91, Group x Shock: F_20,300_=0.7, p=0.74). **D)** No group differences were observed during the trauma recall test (Group: F_2,30_=1.9, p=0.17). **E)** No group differences were observed during the light-dark test of anxiety-related behavior (Group: F_2,30_=1.1, p=0.35). **F)** No group differences were observed during the novel stressor test for stress sensitization (Group: F_2,30_=2.2, p=0.13). mCherry=11, BLA→vHC=11, and vHC→BLA=11 mice. P<.05 (*), p<0.01 (**), p<0.001 (***). Error bars reflect standard error of the mean. Statistics in main text.

## DISCUSSION

Associative learning frameworks – in which cues present at the time of a stressor come to drive behavior – have dominated how we study the impacts of stress on fear and anxiety disorders. Consequently, immense gains have been made in our understanding of the biological basis of associative fear learning, as well how these associations are extinguished. However, relatively little attention has been paid to non-associative learning processes governing stress-induced changes in defensive behavior. Here, utilizing a combination of targeted protein synthesis inhibition, chemogenetic inhibition, and projection-specific inhibition strategies, we demonstrate that the BLA and vHC differentially contribute to stress-induced changes in defensive behavior. Specifically, we find that although both structures contribute to associative learning and recall about a stressful event, they have dissociable contributions to non-associative learning processes: the BLA supports heightened sensitivity to subsequent aversive events, whereas the vHC supports increases in anxiety-related behavior. These findings highlight how both associative and non-associative memories may be formed for a single stressful experience and suggest that independent circuitries support these memories.

A wealth of literature supports the notion that the BLA and vHC regulate defensive behaviors ^43–54^. In light of reciprocal connections between these structures ^77,83,84^, it is often thought that stress-induced plasticity within the BLA and vHC, as well as their coordinated neuronal activity, subserve a common defensive process. While evidence exists that the connectivity of these structures plays an important role in defensive behavior ^79–81,85^, our findings demonstrate that this is not always the case.

First, stress-induced protein synthesis within the BLA was found to be critical to subsequent enhancements in stress sensitivity, whereas stress-induced protein synthesis within the vHC had no bearing on this defensive phenotype. Conversely, stress-induced protein synthesis within the vHC, but not the BLA, was found to support heightened anxiety-related behaviors. Therefore, the fundamental structural plasticity that supports these behavioral changes appears to emerge from distinct brain regions.

Second, it could be the case that neuronal activity within a brain region is necessary to express a particular behavioral change, even when plasticity in that region is not required for that behavioral change to come about. Ruling out this possibility, we found that suppressing neural activity within the BLA and vHC also had doubly dissociable impacts on the expression of these behavioral changes. Inhibiting neural activity in the BLA was able to block the heightened response to aversive stimuli observed after an initial stressor, whereas inhibition of the vHC was without effect. Similarly, inhibiting neural activity in the vHC was able to block stress-induced changes in anxiety-related behavior, whereas inhibition of the BLA was without effect. Accordingly, at both the level of plasticity and neuronal activity, the BLA and vHC appear to differentially regulate these behaviors.

The above findings are surprising given the reciprocal monosynaptic connections between the BLA and vHC. In light of this, we attempted to replicate the effects of BLA/vHC inhibition, this time targeting only the neurons in the BLA that project to the vHC, or only the neurons in the vHC that project to the BLA. Consistent with the hypothesis that the BLA and vHC regulate these behaviors independently, inhibition of either projection was without effect. Therefore, it seems that alternative projections emanating from the BLA and vHC support these behavioral changes. In the future we hope to identify these targets.

The dissociation of the contributions of the BLA and vHC to the defensive behaviors studied here is not entirely without precedent, although side by side comparisons of their functions are limited. For instance, despite the large literature on the role of the BLA in associative fear learning ^9–11,43,44,86–89^, several studies have reported that inhibition of the BLA is without effect on exploratory anxiety-related behaviors ^79,90–92^, although discrepancies exist ^93^. Additionally, a recent report found that optogenetic stimulation of projections from the vHC to the BLA do not regulate exploratory anxiety-related behavior ^80^. This corroborates the hypothesis that vHC regulates exploratory-anxiety related behavior through its connections with other down-stream structures such as the hypothalamus ^80,94,95^ or medial prefrontal cortex ^46^. While stimulation of BLA terminal fibers in the vHC has been found to alter anxiety-related behavior ^79^, this may reflect a general effect of exciting the vHC, as opposed to the natural role served by BLA to vHC efferents. Indeed, we were unable to alter anxiety-related behavior via chemogenetic inhibition of vHC projecting BLA neurons. Together, these results broadly suggest that the vHC may regulate exploratory anxiety-related behavior in a manner distinct from its connections with the BLA.

Our findings also add to existing evidence that associative and non-associative impacts of stress are biologically distinct. Blockade of stress-induced protein synthesis in the BLA and vHC were both able to impair the acquisition of associative fear for the context of the initial stressor, and blockade of activity within either structure was similarly able to impair associative memory recall. Importantly, if the observed changes in anxiety-related behavior and stress sensitivity were dependent upon the associative memory of the stressor, then blocking associative memory for the stressor should impair their expression. However, manipulations of the BLA potently reduced associative memory but did not affect increases in anxiety-related behavior. Likewise, manipulations of the vHC reduced associative memory but did not interfere with stress sensitization. Additionally, it does not seem that these discrepancies are a matter of thresholding (e.g., manipulations of the BLA did not alter the associative memory enough to alter anxiety-related behavior). Blockade of stress-induced protein synthesis in the BLA was more effective at reducing subsequent associative fear of the stressor context than blockade of stress-induced protein synthesis in the vHC. Nevertheless, blocking protein synthesis in the vHC reduced subsequent anxiety-related behavior, whereas the same manipulation in the BLA did not. Therefore, it seems that each of these defensive phenotypes – associative recall, heightened stress sensitivity, and anxiety-related behavior – reflect distinct plasticity mechanisms.

Protein synthesis within the BLA was necessary for the acquisition and expression of both heightened associative memories of a stressful event and the heightened sensitivity to novel stressors after prior stress. Furthermore, these two phenotypes were correlated, albeit weakly. It may be argued that these defensive phenotypes are one in the same. Prior evidence, as well as data presented here, stands in opposition to this possibility. First, extinction of the associative memory for an initial stressor has been found to leave the enhanced response to a second stressor intact ^15,38,39^. Second, early life stress at a time point when rodents are unable to form associative memories nevertheless leaves animals with heightened responses to subsequent stressors in adulthood ^14^. Lastly, despite the finding presented here that inactivation of the BLA and vHC were able to equivalently impair trauma memory recall, only inactivation of the BLA was able to alter the heightened sensitivity to a novel stressor. Therefore, despite both phenotypes’ dependence upon the BLA, associative memory for a stressor and the enhanced responding to subsequent stress are dissociable. It could be that synapse- and ensemble-specific plasticity within the amygdala supports the associative memory for a specific stressor, whereas a broader form of non-associative plasticity within the amygdala supports sensitized stress responses. Future studies will disentangle how plasticity within the amygdala supports these two forms of learning.

In closing, these results shed new light on how stress-induced plasticity within the BLA and vHC support the formation of defensive behavioral phenotypes relevant to neuropsychiatric illness. Furthermore, they highlight just how distinct memories for stressful events might be. Similar to the separate memory systems in the brain supporting episodic and procedural learning of the same event, we find that different defensive behaviors induced by stress are supported by distinct brain regions. This has important clinical implications for the treatment of anxiety disorders and other stress-associated mental health conditions. First, it suggests that clinically targeting one stress-induced defensive behavior, or the circuits that support that behavior, may leave others wholly unaffected. Perhaps some of the existing gaps in treatment result from a failure to adequately target the spectrum of defensive processes altered in these conditions. Second, by understanding the relationship between specific defensive behaviors and their biology, we may make greater headway in the treatment of these conditions. For instance, associative learning processes are more likely affected in some mental health conditions (e.g., PTSD), while anxiety-related behaviors might be more affected in others (e.g., generalized anxiety disorder), and these differences undoubtedly covary with different neuronal patterns. Indeed, it is known that the different anxiety disorders are not only symptomatically different, but characterized by unique brain activity patterns^71^. By understanding these inter-relationships, we may more rapidly find the appropriate key to unlock the door to recovery for stress-associated mental health conditions.

## Supporting information

Key Resources

## ACKNOWLEDGEMENTS

This work was supported by NIMH DP2 MH122399, NIMH R01 MH120162, Brain Research Foundation Award, Klingenstein-Simons Fellowship, NARSAD Young Investigator Award, McKnight Memory and Cognitive Disorder Award, One Mind-Otsuka Rising Star Research Award, Hirschl/Weill-Caulier Award, Mount Sinai Distinguished Scholar Award, and Friedman Brain Institute Award, to DJC; NIMH K99 MH131792 and BBRF Young Investigator Award to ZTP; NINDS F32 NS116416 to ZCW; AES Predoctoral Research Fellowship to YF; and NINDS R01 NS116357 and NIA RF1 AG072497 to TS. The authors would like to thank Dr. Scott Russo and Dr. Roger Clem for their helpful comments on this work.

## AUTHOR CONTRIBUTIONS

ZTP and DJC conceived of the overarching research goals, designed the experiments, and oversaw the experiments. ZTP analyzed the experimental data and prepared the initial manuscript. ZTP, ARL, PS, SDA, BK, ZCW, YF, ZD, MEB, TRF, LC, SLF, IM, TS and DJC contributed to interpretation of the results and edited the manuscript. ZTP, ALB, PS, SDA, BK, ZCW, YF, ZD, MEB, TRF, LC, and SF performed experiments. ZTP and ZD designed software for analysis of behavioral data. DJC, IM, TS, ZTP, ZCW and YF secured funding.

## KEY RESOURCE TABLE

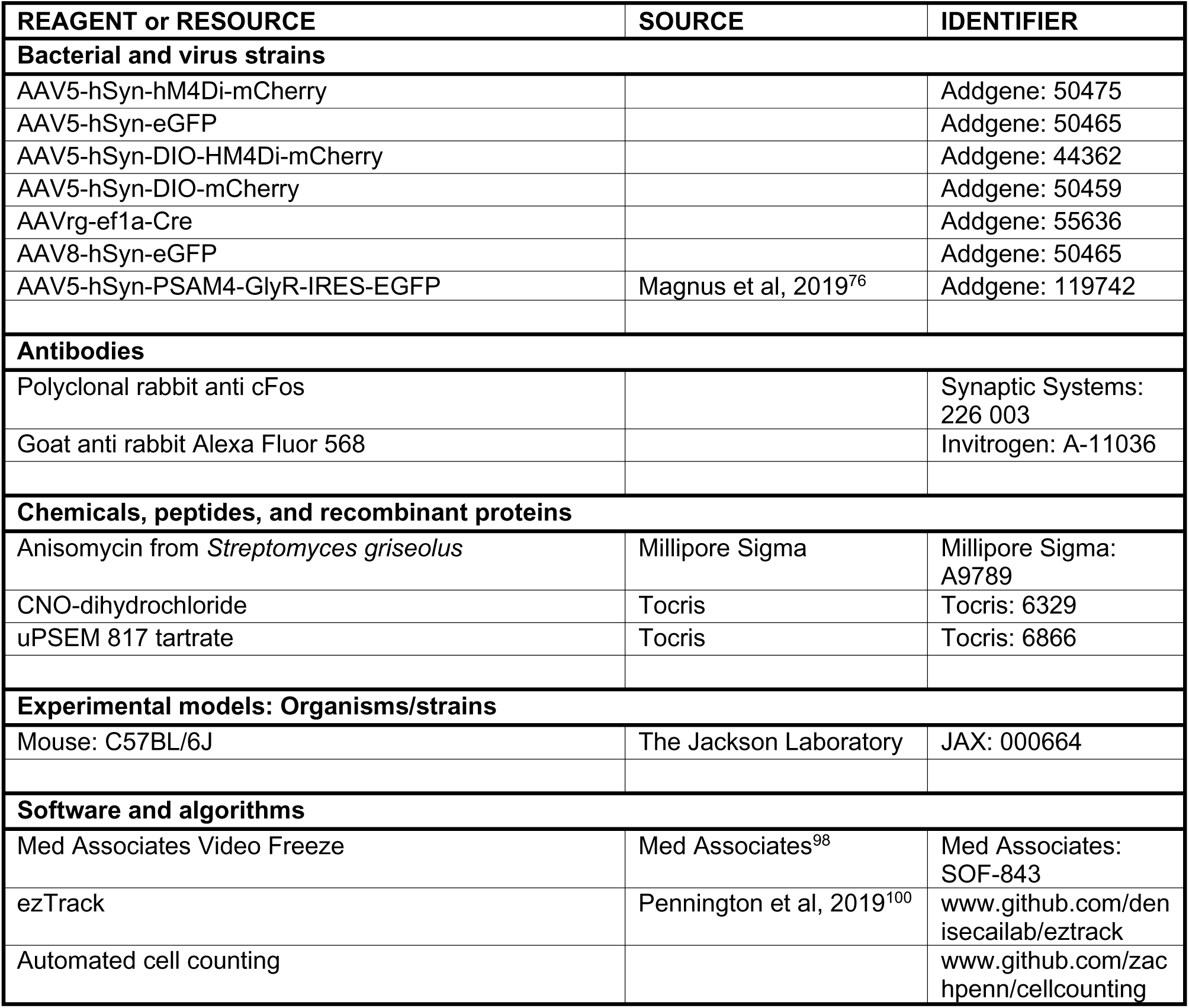

## References and Literature Cited

1. Fendt, M., and Fanselow, M.S. (1999). The neuroanatomical and neurochemical basis of conditioned fear. Neurosci Biobehav Rev 23, 743–760.

2. Bolles, R.C. (1970). Species-specific defense reations and avoidance learning. Psychological Review 77, 32–48.

3. Fanselow, M.S., and Lester, L.S. (1988). A functional behavioristic approach to aversively motivated behavior. In Evolution and Learning, R.C. Bolles, and M.C. Beecher, eds. (Erlbaum), pp. 185–211.

4. Cannon, W.B. (1915). Bodily changes in pain, hunger, fear and rage: An account of recent researches into the function of emotional excitement. (D. Appleton and Componay).

5. Blanchard, R.J., Blanchard, D.C., Rodgers, J., and Weiss, S.M. (1990). The characterization and modelling of antipredator defensive behavior. Neurosci Biobehav Rev 14, 463–472. 10.1016/s0149-7634(05)80069-7.

6. Blanchard, R.J., and Blanchard, D.C. (1969). Passive and active reactions to fear-eliciting stimuli. J Comp Physiol Psychol 68, 129–135. 10.1037/h0027676.

7. Bienvenu, T.C.M., Dejean, C., Jercog, D., Aouizerate, B., Lemoine, M., and Herry, C. (2021). The advent of fear conditioning as an animal model of post-traumatic stress disorder: Learning from the past to shape the future of PTSD research. Neuron 109, 2380–2397. 10.1016/j.neuron.2021.05.017.

8. Maren, S. (2005). Synaptic mechanisms of associative memory in the amygdala. Neuron 47, 783–786. 10.1016/j.neuron.2005.08.009.

9. Maren, S. (2003). The amygdala, synaptic plasticity, and fear memory. Ann N Y Acad Sci 985, 106–113.

10. Fanselow, M.S., and LeDoux, J.E. (1999). Why we think plasticity underlying Pavlovian fear conditioning occurs in the basolateral amygdala. Neuron 23, 229–232.

11. Davis, M. (1992). The role of the amygdala in fear and anxiety. Annu Rev Neurosci 15, 353–375. 10.1146/annurev.ne.15.030192.002033.

12. Fadok, J.P., Krabbe, S., Markovic, M., Courtin, J., Xu, C., Massi, L., Botta, P., Bylund, K., Müller, C., Kovacevic, A., et al. (2017). A competitive inhibitory circuit for selection of active and passive fear responses. Nature 542, 96–100. 10.1038/nature21047.

13. Adamec, R.E., and Shallow, T. (1993). Lasting effects on rodent anxiety of a single exposure to a cat. Physiol Behav 54, 101–109. 10.1016/0031-9384(93)90050-p.

14. Poulos, A.M., Reger, M., Mehta, N., Zhuravka, I., Sterlace, S.S., Gannam, C., Hovda, D.A., Giza, C.C., and Fanselow, M.S. (2014). Amnesia for early life stress does not preclude the adult development of posttraumatic stress disorder symptoms in rats. Biol Psychiatry 76, 306–314. 10.1016/j.biopsych.2013.10.007.

15. Rau, V., DeCola, J.P., and Fanselow, M.S. (2005). Stress-induced enhancement of fear learning: an animal model of posttraumatic stress disorder. Neurosci Biobehav Rev 29, 1207–1223. 10.1016/j.neubiorev.2005.04.010.

16. Pynoos, R.S., Ritzmann, R.F., Steinberg, A.M., Goenjian, A., and Prisecaru, I. (1996). A behavioral animal model of posttraumatic stress disorder featuring repeated exposure to situational reminders. Biol Psychiatry 39, 129–134. 10.1016/0006-3223(95)00088-7.

17. Servatius, R.J., Ottenweller, J.E., and Natelson, B.H. (1995). Delayed startle sensitization distinguishes rats exposed to one or three stress sessions: further evidence toward an animal model of PTSD. Biol Psychiatry 38, 539–546. 10.1016/0006-3223(94)00369-E.

18. American Psychiatric Association. (2013). American Psychiatric Association: Diagnostic and Statistical Manual of Mental Disorders, Fifth Edition. (American Psychiatric Publishing).

19. Moreno-Peral, P., Conejo-Cerón, S., Motrico, E., Rodríguez-Morejón, A., Fernández, A., García-Campayo, J., Roca, M., Serrano-Blanco, A., Rubio-Valera, M., and Bellón, J. (2014). Risk factors for the onset of panic and generalised anxiety disorders in the general adult population: a systematic review of cohort studies. J Affect Disord 168, 337–348. 10.1016/j.jad.2014.06.021.

20. Kessler, R.C., Gruber, M., Hettema, J.M., Hwang, I., Sampson, N., and Yonkers, K.A. (2008). Co-morbid major depression and generalized anxiety disorders in the National Comorbidity Survey follow-up. Psychol Med 38, 365–374. 10.1017/S0033291707002012.

21. Blanco, C., Rubio, J., Wall, M., Wang, S., Jiu, C.J., and Kendler, K.S. (2014). Risk factors for anxiety disorders: common and specific effects in a national sample. Depress Anxiety 31, 756–764. 10.1002/da.22247.

22. Keiser, A.A., Turnbull, L.M., Darian, M.A., Feldman, D.E., Song, I., and Tronson, N.C. (2017). Sex Differences in Context Fear Generalization and Recruitment of Hippocampus and Amygdala during Retrieval. Neuropsychopharmacology 42, 397–407. 10.1038/npp.2016.174.

23. Lissek, S., Bradford, D.E., Alvarez, R.P., Burton, P., Espensen-Sturges, T., Reynolds, R.C., and Grillon, C. (2014). Neural substrates of classically conditioned fear-generalization in humans: a parametric fMRI study. Soc Cogn Affect Neurosci 9, 1134–1142. 10.1093/scan/nst096.

24. Lissek, S., Kaczkurkin, A.N., Rabin, S., Geraci, M., Pine, D.S., and Grillon, C. (2014). Generalized anxiety disorder is associated with overgeneralization of classically conditioned fear. Biol Psychiatry 75, 909–915. 10.1016/j.biopsych.2013.07.025.

25. Zelikowsky, M., Bissiere, S., Hast, T.A., Bennett, R.Z., Abdipranoto, A., Vissel, B., and Fanselow, M.S. (2013). Prefrontal microcircuit underlies contextual learning after hippocampal loss. Proc Natl Acad Sci U S A 110, 9938–9943. 10.1073/pnas.1301691110.

26. Xu, W., and Südhof, T.C. (2013). A neural circuit for memory specificity and generalization. Science 339, 1290–1295. 10.1126/science.1229534.

27. Wiltgen, B.J., Zhou, M., Cai, Y., Balaji, J., Karlsson, M.G., Parivash, S.N., Li, W., and Silva, A.J. (2010). The hippocampus plays a selective role in the retrieval of detailed contextual memories. Curr Biol 20, 1336–1344. 10.1016/j.cub.2010.06.068.

28. Starita, F., Kroes, M.C.W., Davachi, L., Phelps, E.A., and Dunsmoor, J.E. (2019). Threat learning promotes generalization of episodic memory. J Exp Psychol Gen 148, 1426–1434. 10.1037/xge0000551.

29. Dunsmoor, J.E., Kroes, M.C.W., Braren, S.H., and Phelps, E.A. (2017). Threat intensity widens fear generalization gradients. Behav Neurosci 131, 168–175. 10.1037/bne0000186.

30. Dunsmoor, J.E., Otto, A.R., and Phelps, E.A. (2017). Stress promotes generalization of older but not recent threat memories. Proc Natl Acad Sci U S A 114, 9218–9223. 10.1073/pnas.1704428114.

31. Davis, M. (1989). Neural systems involved in fear-potentiated startle. Ann N Y Acad Sci 563, 165–183. 10.1111/j.1749-6632.1989.tb42197.x.

32. Brown, J.S., Kalish, H.I., and Farber, I.E. (1951). Conditioned fear as revealed by magnitude of startle response to an auditory stimulus. J Exp Psychol 41, 317–328. 10.1037/h0060166.

33. Norrholm, S.D., Jovanovic, T., Olin, I.W., Sands, L.A., Karapanou, I., Bradley, B., and Ressler, K.J. (2011). Fear extinction in traumatized civilians with posttraumatic stress disorder: relation to symptom severity. Biol Psychiatry 69, 556–563. 10.1016/j.biopsych.2010.09.013.

34. Jovanovic, T., Blanding, N.Q., Norrholm, S.D., Duncan, E., Bradley, B., and Ressler, K.J. (2009). Childhood abuse is associated with increased startle reactivity in adulthood. Depress Anxiety 26, 1018–1026. 10.1002/da.20599.

35. Kaczkurkin, A.N., Burton, P.C., Chazin, S.M., Manbeck, A.B., Espensen-Sturges, T., Cooper, S.E., Sponheim, S.R., and Lissek, S. (2017). Neural Substrates of Overgeneralized Conditioned Fear in PTSD. Am J Psychiatry 174, 125–134. 10.1176/appi.ajp.2016.15121549.

36. Grillon, C., and Morgan, C.A. (1999). Fear-potentiated startle conditioning to explicit and contextual cues in Gulf War veterans with posttraumatic stress disorder. J Abnorm Psychol 108, 134–142.

37. Acheson, D.T., Geyer, M.A., Baker, D.G., Nievergelt, C.M., Yurgil, K., Risbrough, V.B., and Team, M.-I. (2015). Conditioned fear and extinction learning performance and its association with psychiatric symptoms in active duty Marines. Psychoneuroendocrinology 51, 495–505. 10.1016/j.psyneuen.2014.09.030.

38. Long, V.A., and Fanselow, M.S. (2012). Stress-enhanced fear learning in rats is resistant to the effects of immediate massed extinction. Stress 15, 627–636. 10.3109/10253890.2011.650251.

39. Hassien, A.M., Shue, F., Bernier, B.E., and Drew, M.R. (2020). A mouse model of stress-enhanced fear learning demonstrates extinction-sensitive and extinction-resistant effects of footshock stress. Behav Brain Res 379, 112391. 10.1016/j.bbr.2019.112391.

40. Difede, J., Olden, M., and Cukor, J. (2014). Evidence-based treatment of post-traumatic stress disorder. Annu Rev Med 65, 319–332. 10.1146/annurev-med-051812-145438.

41. Bradley, R., Greene, J., Russ, E., Dutra, L., and Westen, D. (2005). A multidimensional meta-analysis of psychotherapy for PTSD. Am J Psychiatry 162, 214–227. 10.1176/appi.ajp.162.2.214.

42. Cukor, J., Olden, M., Lee, F., and Difede, J. (2010). Evidence-based treatments for PTSD, new directions, and special challenges. Ann N Y Acad Sci 1208, 82–89. 10.1111/j.1749-6632.2010.05793.x.

43. Maren, S., Aharonov, G., and Fanselow, M.S. (1996). Retrograde abolition of conditional fear after excitotoxic lesions in the basolateral amygdala of rats: absence of a temporal gradient. Behav Neurosci 110, 718–726.

44. Blanchard, D.C., and Blanchard, R.J. (1972). Innate and conditioned reactions to threat in rats with amygdaloid lesions. J Comp Physiol Psychol 81, 281–290.

45. Feinstein, J.S., Adolphs, R., Damasio, A., and Tranel, D. (2011). The human amygdala and the induction and experience of fear. Curr Biol 21, 34–38. 10.1016/j.cub.2010.11.042.

46. Padilla-Coreano, N., Bolkan, S.S., Pierce, G.M., Blackman, D.R., Hardin, W.D., Garcia-Garcia, A.L., Spellman, T.J., and Gordon, J.A. (2016). Direct Ventral Hippocampal-Prefrontal Input Is Required for Anxiety-Related Neural Activity and Behavior. Neuron 89, 857–866. 10.1016/j.neuron.2016.01.011.

47. Kheirbek, M.A., Drew, L.J., Burghardt, N.S., Costantini, D.O., Tannenholz, L., Ahmari, S.E., Zeng, H., Fenton, A.A., and Hen, R. (2013). Differential control of learning and anxiety along the dorsoventral axis of the dentate gyrus. Neuron 77, 955–968. 10.1016/j.neuron.2012.12.038.

48. Adhikari, A., Topiwala, M.A., and Gordon, J.A. (2010). Synchronized activity between the ventral hippocampus and the medial prefrontal cortex during anxiety. Neuron 65, 257–269. 10.1016/j.neuron.2009.12.002.

49. Fanselow, M.S., and Dong, H.W. (2010). Are the dorsal and ventral hippocampus functionally distinct structures? Neuron 65, 7–19. 10.1016/j.neuron.2009.11.031.

50. Bannerman, D.M., Yee, B.K., Good, M.A., Heupel, M.J., Iversen, S.D., and Rawlins, J.N. (1999). Double dissociation of function within the hippocampus: a comparison of dorsal, ventral, and complete hippocampal cytotoxic lesions. Behav Neurosci 113, 1170–1188. 10.1037//0735-7044.113.6.1170.

51. Han, J.H., Kushner, S.A., Yiu, A.P., Cole, C.J., Matynia, A., Brown, R.A., Neve, R.L., Guzowski, J.F., Silva, A.J., and Josselyn, S.A. (2007). Neuronal competition and selection during memory formation. Science 316, 457–460. 10.1126/science.1139438.

52. Mastrodonato, A., Martinez, R., Pavlova, I.P., LaGamma, C.T., Brachman, R.A., Robison, A.J., and Denny, C.A. (2018). Ventral CA3 Activation Mediates Prophylactic Ketamine Efficacy Against Stress-Induced Depressive-like Behavior. Biol Psychiatry 84, 846–856. 10.1016/j.biopsych.2018.02.011.

53. Zaki, Y., Mau, W., Cincotta, C., Monasterio, A., Odom, E., Doucette, E., Grella, S.L., Merfeld, E., Shpokayte, M., and Ramirez, S. (2022). Hippocampus and amygdala fear memory engrams re-emerge after contextual fear relapse. Neuropsychopharmacology 47, 1992–2001. 10.1038/s41386-022-01407-0.

54. Ramirez, S., Liu, X., MacDonald, C.J., Moffa, A., Zhou, J., Redondo, R.L., and Tonegawa, S. (2015). Activating positive memory engrams suppresses depression-like behaviour. Nature 522, 335–339. 10.1038/nature14514.

55. Stevens, J.S., Almli, L.M., Fani, N., Gutman, D.A., Bradley, B., Norrholm, S.D., Reiser, E., Ely, T.D., Dhanani, R., Glover, E.M., et al. (2014). PACAP receptor gene polymorphism impacts fear responses in the amygdala and hippocampus. Proc Natl Acad Sci U S A 111, 3158–3163. 10.1073/pnas.1318954111.

56. Perusini, J., Meyer, E., Rau, V., Avershal, J., Rajbhandari, A., Hoffman, A., Nocera, N., Condro, M., Waschek, J., Spigelman, I., and Fanselow, M. (2014). Mechanisms underlying the induction and expression of fear sensitization following acute traumatic stress. Pavlovian Society.

57. Rau, V., and Fanselow, M.S. (2009). Exposure to a stressor produces a long lasting enhancement of fear learning in rats. Stress 12, 125–133. 10.1080/10253890802137320.

58. Pennington, Z.T., Trott, J.M., Rajbhandari, A.K., Li, K., Walwyn, W.M., Evans, C.J., and Fanselow, M.S. (2020). Chronic opioid pretreatment potentiates the sensitization of fear learning by trauma. Neuropsychopharmacology 45, 482–490. 10.1038/s41386-019-0559-5.

59. Chaouloff, F., Durand, M., and Mormède, P. (1997). Anxiety- and activity-related effects of diazepam and chlordiazepoxide in the rat light/dark and dark/light tests. Behav Brain Res 85, 27–35. 10.1016/s0166-4328(96)00160-x.

60. Crawley, J.N. (1981). Neuropharmacologic specificity of a simple animal model for the behavioral actions of benzodiazepines. Pharmacol Biochem Behav 15, 695–699. 10.1016/0091-3057(81)90007-1.

61. Kessler, R.C., Sonnega, A., Bromet, E., Hughes, M., and Nelson, C.B. (1995). Posttraumatic stress disorder in the National Comorbidity Survey. Arch Gen Psychiatry 52, 1048–1060.

62. Breslau, N., Davis, G.C., Andreski, P., Peterson, E.L., and Schultz, L.R. (1997). Sex differences in posttraumatic stress disorder. Arch Gen Psychiatry 54, 1044–1048.

63. Smith, A.C.W., Jonkman, S., Difeliceantonio, A.G., O’Connor, R.M., Ghoshal, S., Romano, M.F., Everitt, B.J., and Kenny, P.J. (2021). Opposing roles for striatonigral and striatopallidal neurons in dorsolateral striatum in consolidating new instrumental actions. Nat Commun 12, 5121. 10.1038/s41467-021-25460-3.

64. Hernandez, P.J., Sadeghian, K., and Kelley, A.E. (2002). Early consolidation of instrumental learning requires protein synthesis in the nucleus accumbens. Nat Neurosci 5, 1327–1331. 10.1038/nn973.

65. Santini, E., Ge, H., Ren, K., Peña de Ortiz, S., and Quirk, G.J. (2004). Consolidation of fear extinction requires protein synthesis in the medial prefrontal cortex. J Neurosci 24, 5704–5710. 10.1523/JNEUROSCI.0786-04.2004.

66. Schafe, G.E., and LeDoux, J.E. (2000). Memory consolidation of auditory pavlovian fear conditioning requires protein synthesis and protein kinase A in the amygdala. J Neurosci 20, RC96.

67. Nader, K., Schafe, G.E., and Le Doux, J.E. (2000). Fear memories require protein synthesis in the amygdala for reconsolidation after retrieval. Nature 406, 722–726. 10.1038/35021052.

68. Bourtchouladze, R., Abel, T., Berman, N., Gordon, R., Lapidus, K., and Kandel, E.R. (1998). Different training procedures recruit either one or two critical periods for contextual memory consolidation, each of which requires protein synthesis and PKA. Learn Mem 5, 365–374.

69. Kandel, E.R. (2001). The molecular biology of memory storage: a dialogue between genes and synapses. Science 294, 1030–1038. 10.1126/science.1067020.

70. Gilbertson, M.W., Shenton, M.E., Ciszewski, A., Kasai, K., Lasko, N.B., Orr, S.P., and Pitman, R.K. (2002). Smaller hippocampal volume predicts pathologic vulnerability to psychological trauma. Nat Neurosci 5, 1242–1247. 10.1038/nn958.

71. Etkin, A., and Wager, T.D. (2007). Functional neuroimaging of anxiety: a meta-analysis of emotional processing in PTSD, social anxiety disorder, and specific phobia. Am J Psychiatry 164, 1476–1488. 10.1176/appi.ajp.2007.07030504.

72. Kühn, S., and Gallinat, J. (2013). Gray matter correlates of posttraumatic stress disorder: a quantitative meta-analysis. Biol Psychiatry 73, 70–74. 10.1016/j.biopsych.2012.06.029.

73. Rabinak, C.A., Angstadt, M., Welsh, R.C., Kenndy, A.E., Lyubkin, M., Martis, B., and Phan, K.L. (2011). Altered amygdala resting-state functional connectivity in post-traumatic stress disorder. Front Psychiatry 2, 62. 10.3389/fpsyt.2011.00062.

74. Armbruster, B.N., Li, X., Pausch, M.H., Herlitze, S., and Roth, B.L. (2007). Evolving the lock to fit the key to create a family of G protein-coupled receptors potently activated by an inert ligand. Proc Natl Acad Sci U S A 104, 5163–5168. 10.1073/pnas.0700293104.

75. Sierra-Mercado, D., Padilla-Coreano, N., and Quirk, G.J. (2011). Dissociable roles of prelimbic and infralimbic cortices, ventral hippocampus, and basolateral amygdala in the expression and extinction of conditioned fear. Neuropsychopharmacology 36, 529–538. 10.1038/npp.2010.184.

76. Magnus, C.J., Lee, P.H., Bonaventura, J., Zemla, R., Gomez, J.L., Ramirez, M.H., Hu, X., Galvan, A., Basu, J., Michaelides, M., and Sternson, S.M. (2019). Ultrapotent chemogenetics for research and potential clinical applications. Science 364. 10.1126/science.aav5282.

77. Hintiryan, H., Bowman, I., Johnson, D.L., Korobkova, L., Zhu, M., Khanjani, N., Gou, L., Gao, L., Yamashita, S., Bienkowski, M.S., et al. (2021). Connectivity characterization of the mouse basolateral amygdalar complex. Nat Commun 12, 2859. 10.1038/s41467-021-22915-5.

78. Gergues, M.M., Han, K.J., Choi, H.S., Brown, B., Clausing, K.J., Turner, V.S., Vainchtein, I.D., Molofsky, A.V., and Kheirbek, M.A. (2020). Circuit and molecular architecture of a ventral hippocampal network. Nat Neurosci 23, 1444–1452. 10.1038/s41593-020-0705-8.

79. Felix-Ortiz, A.C., Beyeler, A., Seo, C., Leppla, C.A., Wildes, C.P., and Tye, K.M. (2013). BLA to vHPC inputs modulate anxiety-related behaviors. Neuron 79, 658–664. 10.1016/j.neuron.2013.06.016.

80. Jimenez, J.C., Su, K., Goldberg, A.R., Luna, V.M., Biane, J.S., Ordek, G., Zhou, P., Ong, S.K., Wright, M.A., Zweifel, L., et al. (2018). Anxiety Cells in a Hippocampal-Hypothalamic Circuit. Neuron 97, 670–683.e676. 10.1016/j.neuron.2018.01.016.

81. Jimenez, J.C., Berry, J.E., Lim, S.C., Ong, S.K., Kheirbek, M.A., and Hen, R. (2020). Contextual fear memory retrieval by correlated ensembles of ventral CA1 neurons. Nat Commun 11, 3492. 10.1038/s41467-020-17270-w.

82. Beyeler, A., Namburi, P., Glober, G.F., Simonnet, C., Calhoon, G.G., Conyers, G.F., Luck, R., Wildes, C.P., and Tye, K.M. (2016). Divergent Routing of Positive and Negative Information from the Amygdala during Memory Retrieval. Neuron 90, 348–361. 10.1016/j.neuron.2016.03.004.

83. Canteras, N.S., Simerly, R.B., and Swanson, L.W. (1992). Connections of the posterior nucleus of the amygdala. J Comp Neurol 324, 143–179. 10.1002/cne.903240203.

84. Canteras, N.S., and Swanson, L.W. (1992). Projections of the ventral subiculum to the amygdala, septum, and hypothalamus: a PHAL anterograde tract-tracing study in the rat. J Comp Neurol 324, 180–194. 10.1002/cne.903240204.

85. Jackson, A.D., Cohen, J.L., Phensy, A.J., Chang, E.F., Dawes, H.E., and Sohal, V.S. (2024). Amygdala-hippocampus somatostatin interneuron beta-synchrony underlies a cross-species biomarker of emotional state. Neuron. 10.1016/j.neuron.2023.12.017.

86. Sprengelmeyer, R., Young, A.W., Schroeder, U., Grossenbacher, P.G., Federlein, J., Büttner, T., and Przuntek, H. (1999). Knowing no fear. Proc Biol Sci 266, 2451–2456. 10.1098/rspb.1999.0945.

87. Fanselow, M.S., and Kim, J.J. (1994). Acquisition of contextual Pavlovian fear conditioning is blocked by application of an NMDA receptor antagonist D,L-2-amino-5-phosphonovaleric acid to the basolateral amygdala. Behav Neurosci 108, 210–212.

88. Rumpel, S., LeDoux, J., Zador, A., and Malinow, R. (2005). Postsynaptic receptor trafficking underlying a form of associative learning. Science 308, 83–88. 10.1126/science.1103944.

89. Han, J.H., Kushner, S.A., Yiu, A.P., Hsiang, H.L., Buch, T., Waisman, A., Bontempi, B., Neve, R.L., Frankland, P.W., and Josselyn, S.A. (2009). Selective erasure of a fear memory. Science 323, 1492–1496. 10.1126/science.1164139.

90. Ribeiro, A.M., Barbosa, F.F., Munguba, H., Costa, M.S., Cavalcante, J.S., and Silva, R.H. (2011). Basolateral amygdala inactivation impairs learned (but not innate) fear response in rats. Neurobiol Learn Mem 95, 433–440. 10.1016/j.nlm.2011.02.004.

91. Tye, K.M., Prakash, R., Kim, S.Y., Fenno, L.E., Grosenick, L., Zarabi, H., Thompson, K.R., Gradinaru, V., Ramakrishnan, C., and Deisseroth, K. (2011). Amygdala circuitry mediating reversible and bidirectional control of anxiety. Nature 471, 358–362. 10.1038/nature09820.

92. Moreira, C.M., Masson, S., Carvalho, M.C., and Brandão, M.L. (2007). Exploratory behaviour of rats in the elevated plus-maze is differentially sensitive to inactivation of the basolateral and central amygdaloid nuclei. Brain Res Bull 71, 466–474. 10.1016/j.brainresbull.2006.10.004.

93. Bueno, C.H., Zangrossi, H., and Viana, M.B. (2005). The inactivation of the basolateral nucleus of the rat amygdala has an anxiolytic effect in the elevated T-maze and light/dark transition tests. Braz J Med Biol Res 38, 1697–1701. 10.1590/s0100-879x2005001100019.

94. Bang, J.Y., Sunstrum, J.K., Garand, D., Parfitt, G.M., Woodin, M., Inoue, W., and Kim, J. (2022). Hippocampal-hypothalamic circuit controls context-dependent innate defensive responses. Elife 11. 10.7554/eLife.74736.

95. Yan, J.J., Ding, X.J., He, T., Chen, A.X., Zhang, W., Yu, Z.X., Cheng, X.Y., Wei, C.Y., Hu, Q.D., Liu, X.Y., et al. (2022). A circuit from the ventral subiculum to anterior hypothalamic nucleus GABAergic neurons essential for anxiety-like behavioral avoidance. Nat Commun 13, 7464. 10.1038/s41467-022-35211-7.

96. Ozawa, T., Ycu, E.A., Kumar, A., Yeh, L.F., Ahmed, T., Koivumaa, J., and Johansen, J.P. (2017). A feedback neural circuit for calibrating aversive memory strength. Nat Neurosci 20, 90–97. 10.1038/nn.4439.

97. Rescorla, R.W., and Wagner, A.R. (1972). A theory of Pavlovian conditioning: Variations in the effectiveness of reinforcement and nonreinforcement. In Classical conditioning: II. Current research and theory, B.A. H, and P.W. F, eds. (Appleton-Century-Crofts), pp. 64–99.

98. Anagnostaras, S.G., Wood, S.C., Shuman, T., Cai, D.J., Leduc, A.D., Zurn, K.R., Zurn, J.B., Sage, J.R., and Herrera, G.M. (2010). Automated assessment of pavlovian conditioned freezing and shock reactivity in mice using the video freeze system. Front Behav Neurosci 4. 10.3389/fnbeh.2010.00158.

99. Pennington, Z.T., Diego, K.S., Francisco, T.R., LaBanca, A.R., Lamsifer, S.I., Liobimova, O., Shuman, T., and Cai, D.J. (2021). ezTrack-A Step-by-Step Guide to Behavior Tracking. Curr Protoc 1, e255. 10.1002/cpz1.255.

100. Pennington, Z.T., Dong, Z., Feng, Y., Vetere, L.M., Page-Harley, L., Shuman, T., and Cai, D.J. (2019). ezTrack: An open-source video analysis pipeline for the investigation of animal behavior. Sci Rep 9, 19979. 10.1038/s41598-019-56408-9.

101. Lattal, K.M., and Abel, T. (2001). Different requirements for protein synthesis in acquisition and extinction of spatial preferences and context-evoked fear. J Neurosci 21, 5773–5780. 10.1523/JNEUROSCI.21-15-05773.2001.

102. Frankland, P.W., Ding, H.K., Takahashi, E., Suzuki, A., Kida, S., and Silva, A.J. (2006). Stability of recent and remote contextual fear memory. Learn Mem 13, 451–457. 10.1101/lm.183406.

103. Lattal, K.M., and Abel, T. (2004). Behavioral impairments caused by injections of the protein synthesis inhibitor anisomycin after contextual retrieval reverse with time. Proc Natl Acad Sci U S A 101, 4667–4672. 10.1073/pnas.0306546101.

104. Grecksch, G., and Matthies, H. (1980). Two sensitive periods for the amnesic effect of anisomycin. Pharmacol Biochem Behav 12, 663–665. 10.1016/0091-3057(80)90145-8.

105. Wanisch, K., and Wotjak, C.T. (2008). Time course and efficiency of protein synthesis inhibition following intracerebral and systemic anisomycin treatment. Neurobiol Learn Mem 90, 485–494. 10.1016/j.nlm.2008.02.007.

106. Flood, J.F., Rosenzweig, M.R., Bennett, E.L., and Orme, A.E. (1973). The influence of duration of protein synthesis inhibition on memory. Physiol Behav 10, 555–562. 10.1016/0031-9384(73)90221-7.

107. Franklin, K., and Paxinos, G. (2008). The mouse brain in stereotaxic coordinates, 3 Edition (Elsevier Inc.).

108. Ko, B., Yoo, J.Y., Yoo, T., Choi, W., Dogan, R., Sung, K., Um, D., Lee, S.B., Kim, H.J., Lee, S., et al. (2023). Npas4-mediated dopaminergic regulation of safety memory consolidation. Cell Rep 42, 112678. 10.1016/j.celrep.2023.112678.

