## Supplemental Figures for "Dissociable contributions of the amygdala and ventral hippocampus to stress-induced changes in defensive behavior"

### TABLE OF CONTENTS

**Supplemental Figures ………………………… 3-16**

**Supplemental Methods ………………………... 17-24**

**References ……………………………………… 25**

### SUPPLEMENTAL FIGURES

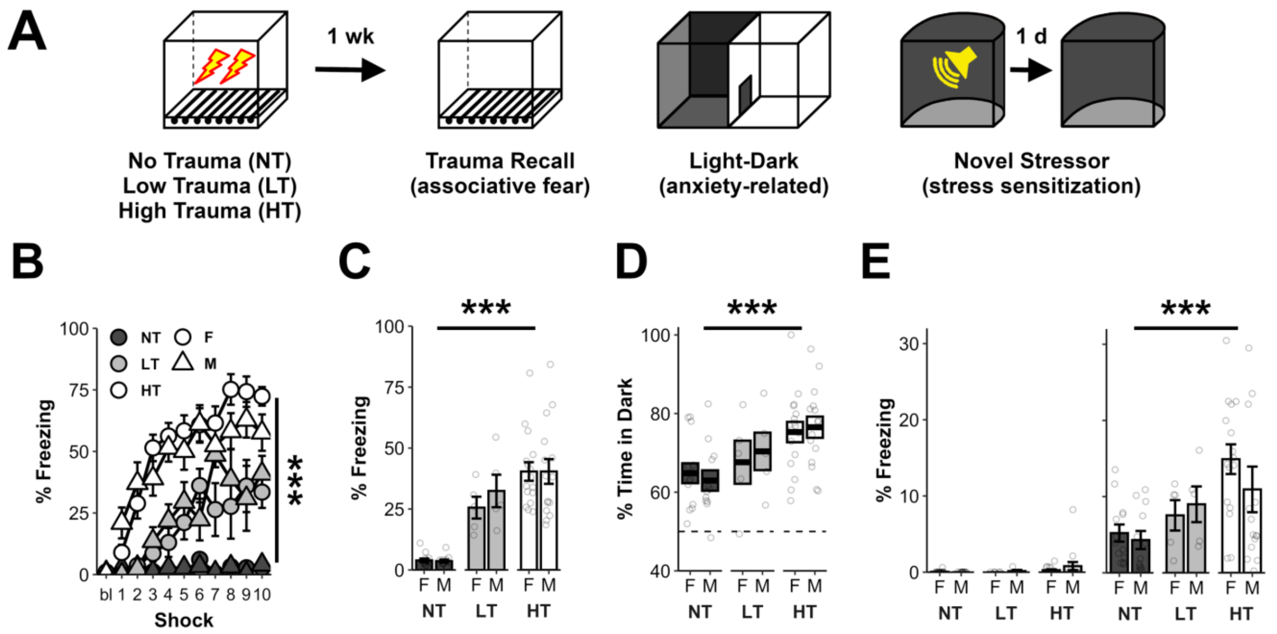


**Figure S1: Trauma-induced changes in defensive behavior are dependent upon trauma-severity, but not sex.**

**A)** Animals were placed in a discrete environment and received either a low amplitude trauma (LT. 10, 0.25 mA shocks); a high amplitude trauma (HT. 10, 1 mA shocks); or were placed in the same environment but were not shocked (NT). A week later, they were tested for associative fear of the trauma environment, anxiety-related behavior in the light-dark test, and stress sensitization, by exposing them to a novel stressor in a new environment. These are the same animals as in Fig 1, here including the ‘T: Low’ group.

**B)** Trauma-exposed animals displayed high levels of post-shock freezing during the trauma, proportional to shock strength (Group: $F_{2,60}$=136.9, p<0.001; Group X Shock: $F_{20,600}$=14, p<0.001). No effects of sex were observed (Sex: $F_{1,60}$=0, p=0.91; Group X Sex: $F_{2,60}$=1, p=0.39; Group X Sex X Shock: $F_{20,600}$=1.3, p=0.23).

**C)** Associative fear was proportional to trauma strength (Group: $F_{2,60}$=75.3, p<0.001). No effects of sex were observed (Sex: $F_{1,60}$=0.3, p=0.56; Group X Sex: $F_{2,60}$=0.3, p=0.73).

**D)** Anxiety-related behavior in the light-dark test was proportional to trauma strength (Group: $F_{2,60}$=9.9, p<0.001). No effects of sex were observed (Sex: $F_{1,60}$=0, p=0.83; Group X Sex: $F_{2,60}$=0.2, p=0.8).

**E)** No differences were observed between groups when examining freezing at baseline prior to the novel stressor (left panel. Group: $F_{2,60}$=1.7, p=0.19). However, trauma-exposed animals showed heightened freezing on the final test day, proportional to trauma strength (right panel. Group: $F_{2,60}$=8.7, p<0.001). No effects of sex were observed at baseline (Sex: $F_{1,60}$=0.5, p=0.48; Group X Sex: $F_{2,60}$=0.5, p=0.6), nor in the final test (Sex: $F_{1,60}$=0.4, p=0.52; Group X Sex: $F_{2,60}$=0.6, p=0.56).

NT=25(13 female), LT=10(5 female), and HT=31(16 female) mice. p<.05 (*), p<0.01 (**), p<0.001 (***). Error bars reflect standard error of the mean.

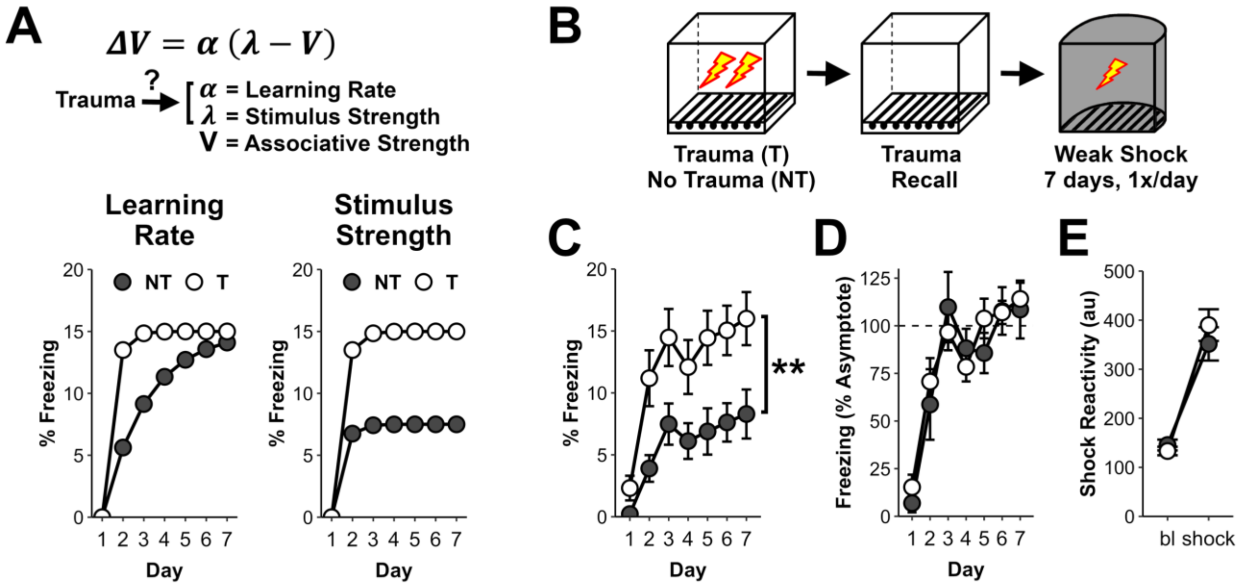


**Figure S2: Trauma impacts learning consistent with changes in stimulus sensitivity.**

**A)** Prior reports indicate that stress augments the learning of new associative fear responses [1]. However, according to theoretical accounts that model trial-by-trial learning such as the Rescorla-Wagner model (top) [2], prior stress could influence associative fear learning in at least two ways. First, prior stress could lead to heightened responding to stimuli (CS) paired with a stressor (US) by altering the rate of learning. If true, with repeated CS-US pairings trauma-exposed (T) and control animals (NT) would come to display similar levels of freezing in response to the CS (bottom left). Alternatively, it could be that trauma-exposed animals perceive the US as stronger (i.e., enhanced sensitivity). In this case, trauma-exposed animals are expected to continually freeze more (bottom right). Of note, asymptotic freezing levels have been confirmed to be proportional to US strength [3].

**B)** To test these alternatives, animals experienced a 10 shock trauma, or were placed in the same environment but received no shocks. The next day, animals were placed back in the trauma environment for a recall session (NT=8.5% vs T=60.5% freezing). Lastly, all animals were placed in a second environment and received one weak shock per day for 7 days.

**C)** Across the course of 7 days of weak shock conditioning, trauma-exposed animals froze significantly more than controls, consistent with inflated stimulus strength (Trauma: $F_{1,22}$=10.2, p<0.01; Trauma X Day: $F_{6,132}$=1.4, p=0.21).

**D)** To examine if learning rate differences might also be present, we normalized each mouse's daily freezing to their asymptotic level of freezing (defined as average freezing across days 3-7). Here, we found no differences between groups (Trauma: $F_{1,22}$=0.8, p=0.39; Trauma X Day: $F_{6,132}$=0.4, p=0.77). These findings indicate that stimulus strength, but not learning rate, is altered by the stressor used here.

**E)** To assess if sensitivity to stress differed at the level of reflexive sensory-motor changes, we examined shock reactivity (motion before and during the 2 sec weak shock). Here, no differences in shock reactivity were observed (Trauma: $F_{1,22}$=0.3, p=0.62; Trauma x Time: $F_{1,22}$=1.2, p=0.28; Time: $F_{1,22}$=102, p<0.001). This suggests that the sensitization likely reflects heightened internal valence, as opposed to more peripheral sensory-motor sensitivity.

NT=12 and T=12 mice. p<.05 (*), p<0.01 (**), p<0.001 (***). Error bars reflect standard error of the mean. au=arbitrary units

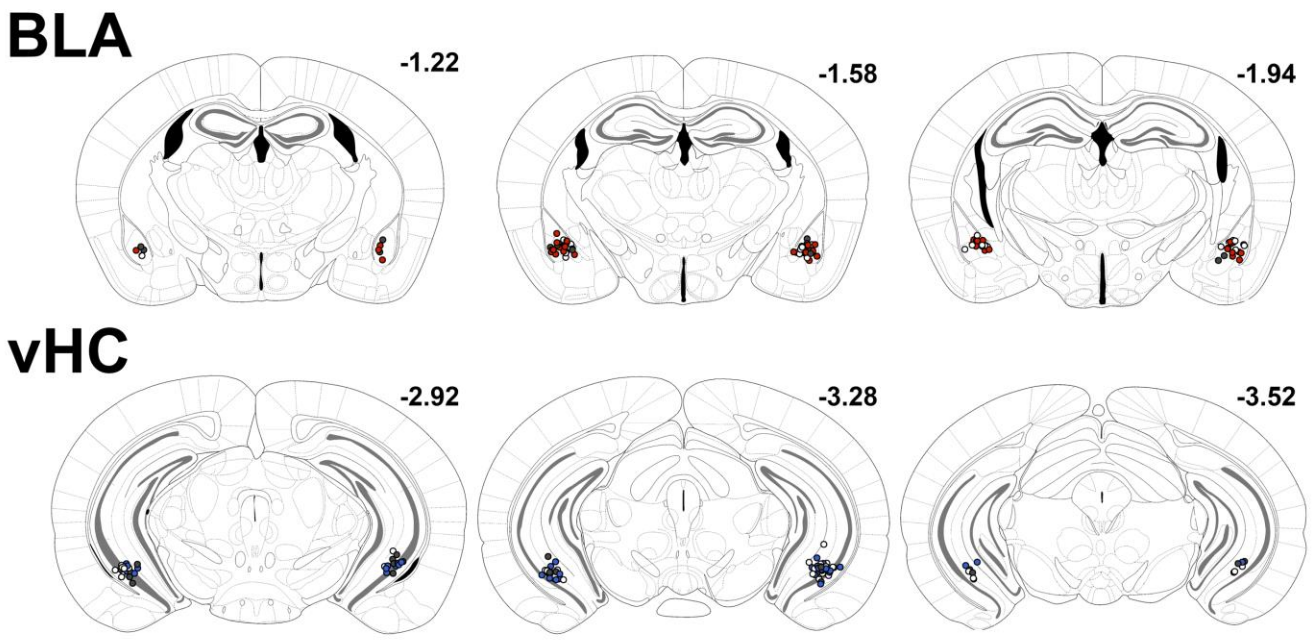


**Figure S3: Placement of injector tips for cannulation of BLA/vHC in main Figure 2.**

For BLA, all cannula tips were located within the BLA, with some bordering the basomedial amygdala. For vHC, all cannula tips were located in ventral CA1. Colors correspond to groups in the main figure (NT: veh=gray; T: veh=white; T: ani-BLA=red; T: ani-vHC=blue). Numbers adjacent to each coronal section correspond to anterior-posterior distance from bregma, in mm, according to the atlas of Franklin and Paxinos [4].

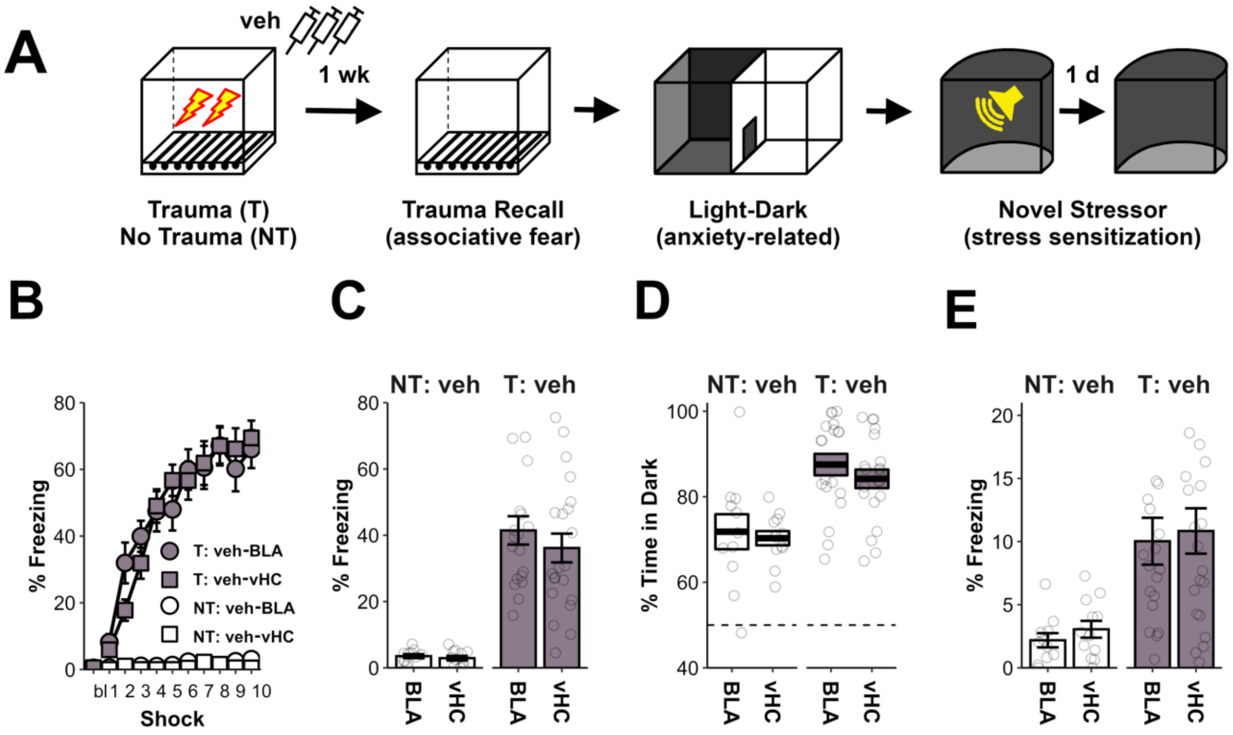


**Figure S4: No differences in animals receiving cannula infusions of vehicle in BLA versus vHC.**

**A)** For the experiment presented in Figure 2F-J, animals with cannula in BLA and vHC that received vehicle infusions are collapsed (NT: veh and T: veh). Here, data from vehicle treated animals is provided in full to demonstrate an absence of an effect of cannula/infusion location.

**B)** There were no differences in freezing during the trauma session (Region: $F_{1,59}$=0, p=0.85; Region X Trauma: $F_{1,59}$=0, p=0.99; Region X Shock: $F_{110,590}$=0.6, p=0.75; Region X Trauma X Shock: $F_{110,590}$=0.8, p=0.56).

**C)** There were no differences in freezing in the trauma recall session (Region: $F_{1,59}$=0.9, p=0.35; Region X Trauma: $F_{1,59}$=0.6, p=0.46).

**D)** There were no differences in time spent in the dark side in the light-dark test (Region: $F_{1,59}$=0.7, p=0.4; Region X Trauma: $F_{1,59}$=0.1, p=0.75).

**E)** There were no differences in freezing in the final novel stressor test session (Region: $F_{1,59}$=0.4, p=0.55; Region X Trauma: $F_{1,59}$=0, p=0.98).

T: veh-BLA=19, T: veh-VHC=21, NT: veh-BLA=11 and NT: veh-vHC=12 mice. p<.05 (*), p<0.01 (**), p<0.001 (***). Error bars reflect standard error of the mean.

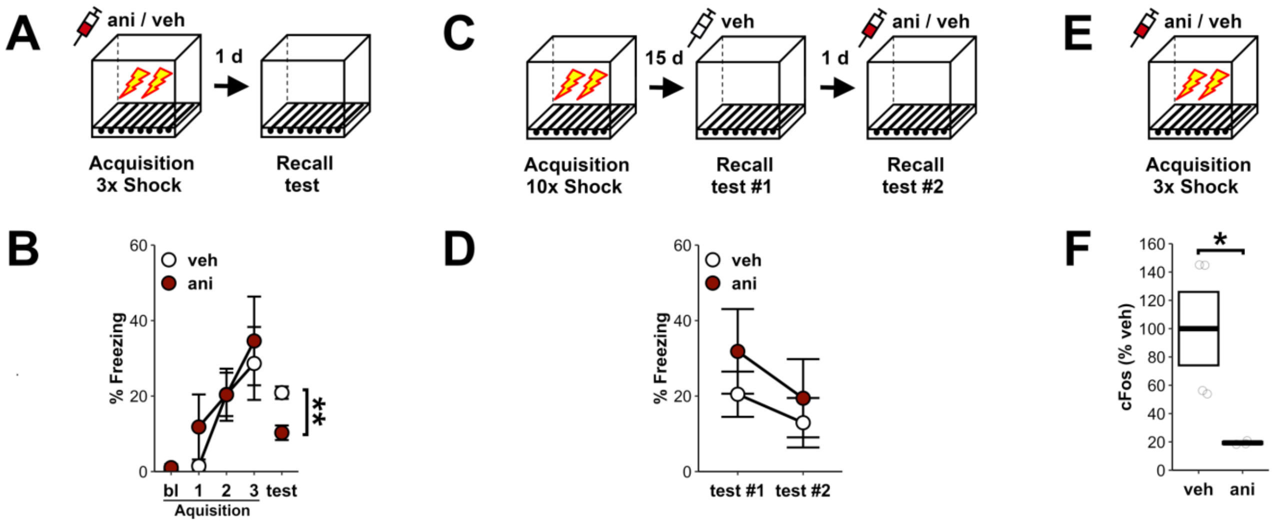


**Figure S5: Anisomycin alters learning, but not expression, of associative fear.**

Reports indicate that anisomycin at high doses can produce off-target effects, including the suppression of neuronal activity [5]. To validate that the concentration of anisomycin (ani) used was not producing effects on behavior that would be consistent with these off-target effects, we assessed the impact of infusing ani into the BLA on fear memory encoding and retention. Notably, we used a concentration that was substantially lower than most prior reports (10 mg/ml here vs >100 mg/ml [6-8]).

**A)** In a first experiment (A-B), we infused vehicle (veh) or ani (10 mg/ml) into the BLA 20 minutes prior to conditioning mice with 3, 1 sec, 1mA, shocks. The next day, we returned the mice to the conditioning context to assess associative fear recall.

**B)** During the conditioning session, animals treated with ani and veh did not show differences in freezing, either prior to the first shock at baseline (bl; $F_{1,7}$=0.3, p=0.62) or after each of the 3 administered shocks (Group: $F_{1,7}$=0.3, p=0.58; Group x Shock: $F_{2,14}$=0.3, p=0.72). Additionally, there was an increase in freezing across the course of the acquisition session, reflecting intact short-term memory (Shock: $F_{2,14}$=7.6, p<0.01). This suggests that ani was not having acute effects on neuronal activity, as disruption of neuronal activity within the amygdala is expected to decrease freezing. That said, ani was able to impair memory consolidation, demonstrated by reduced freezing when animals treated with ani were returned to the context for a recall test the next day (Group: $F_{1,7}$=13.6, p<0.01).

**C)** In a second experiment, we gave animals 10, 1 sec, 1 mA, shocks over the course of an hour (Trauma). We subsequently examined freezing when animals were returned to trauma context during two recall tests. In the first test, all animals were given veh 20 min prior to a 5 min recall. In the second test, half of the animals were given veh and the other half of animals were given ani.

**D)** Notably, there were no acute effects of ani on freezing behavior at recall, further confirming that the dose of ani used does not acutely alter amygdala dependent behavior (Group: $F_{1,7}$: 0.48, p=0.51; Group x Test: $F_{1,7}$=0.44, p=0.53).

**E)** Animals were given either veh/ani and conditioned 30 minutes later to induce cFos. 60 minutes after behavioral testing, tissue was collected for cFos immunohistochemistry.

**F)** cFos translation was significantly reduced by anisomycin, by approximately 80%, evidence that the dose used effectively blocked protein synthesis ($F_{1,5}$=7.3, p=0.04).

For effect of ani on acquisition in A-B: veh=4, ani=5. For effect of ani on recall in C-D: veh=8, ani=10. For effect of ani on cFos in C-D: veh=4, ani=3. p<.05 (*), p<0.01 (**), p<0.001 (***). Error bars reflect standard error of the mean.

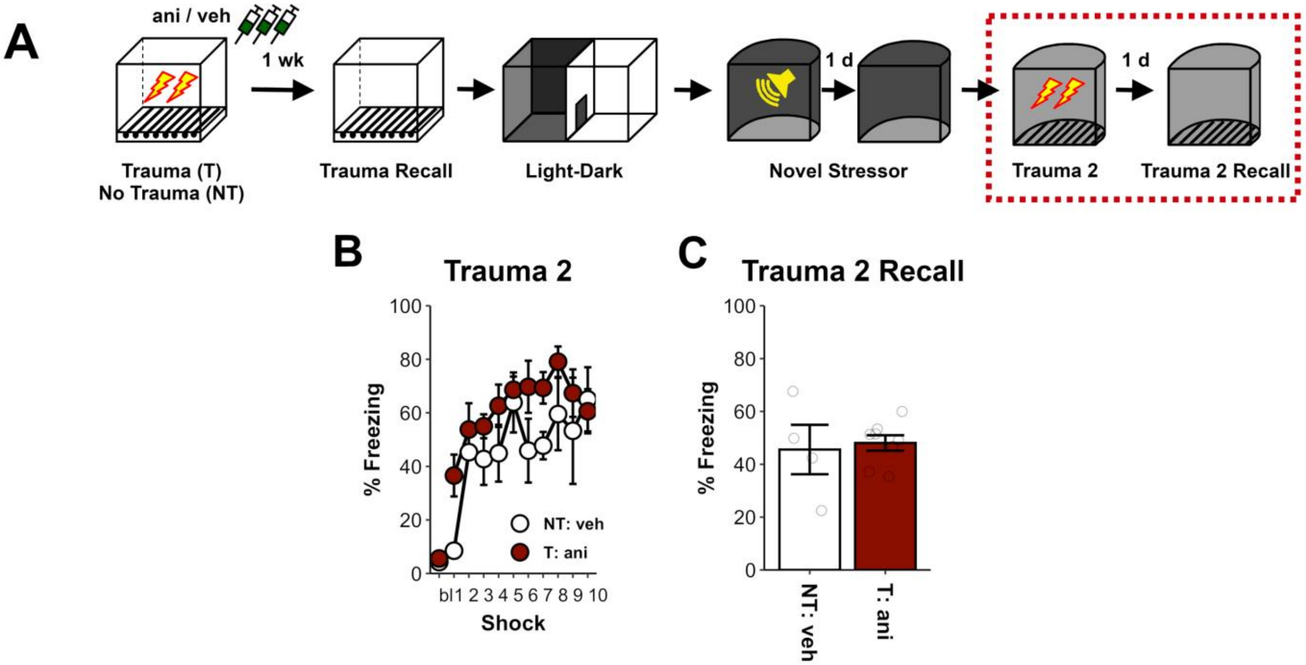


**Figure S6: Prior anisomycin in the amygdala does not permanently impair the ability to form associative fear memories.**

**A)** In order to confirm that repeated anisomycin injections do not produce permanent impairments in associative learning, a subset of mice that had previously received trauma with anisomycin in the amygdala (T: ani) were given a second trauma in a novel context (Fig 2 in Main Text). They were compared to animals that previously received no trauma and vehicle in the BLA (NT: veh).

**B)** During the second trauma, the two groups showed no difference in baseline freezing (bl; $F_{1,10}$=0.2, p=0.64). Moreover, there were no differences in post-shock freezing (Group: $F_{1,10}$=4.26, p=0.07; Group X Shock: $F_{9,90}$=0.7, p=0.73). However, freezing increased across the session, evidence of learning (Shock: $F_{9,90}$=5, p<0.01).

**C)** In a recall test, there were no group differences in freezing ($F_{1,10}$=0.1, p=0.83). These data suggest that administration of anisomycin did not produce permanent alterations to the amygdala, as associative freezing is profoundly impaired by amygdala damage [9-12].

NT: veh=4, T: ani=8. p<.05 (*), p<0.01 (**), p<0.001 (***). Error bars reflect standard error of the mean.

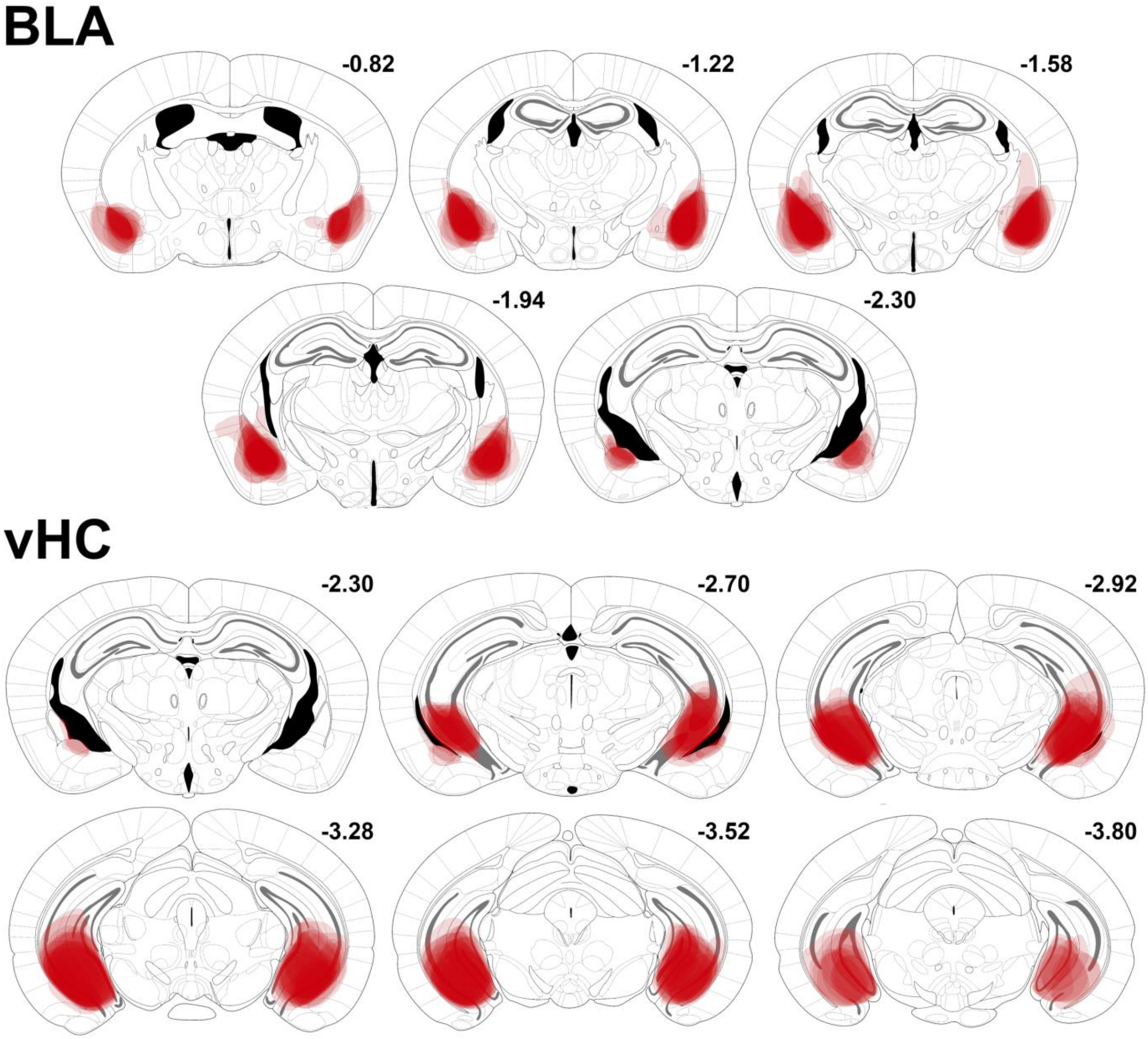


**Figure S7: Distribution of HM4D expression in BLA/vHC.**

Here, expression of HM4D is depicted for main figure 3F-J. Distribution of viral expression of HM4D followed the same pattern across experiments. For BLA injections, viral expression was largely focused in the BLA and lateral amygdala, with moderate spread to central and basomedial amygdala compartments. Light spread was also occasionally observed in adjacent piriform cortex. For the vHC, viral expression was most heavily focused in ventral CA1 and subiculum, with moderate expression in CA3 and dentate gyrus. Light spread was also occasionally observed in adjacent entorhinal cortex and most posterior BLA. Spread for every included animal is displayed. Numbers adjacent to each coronal section correspond to anterior-posterior distance from bregma, in mm, according to the atlas of Franklin and Paxinos [4].

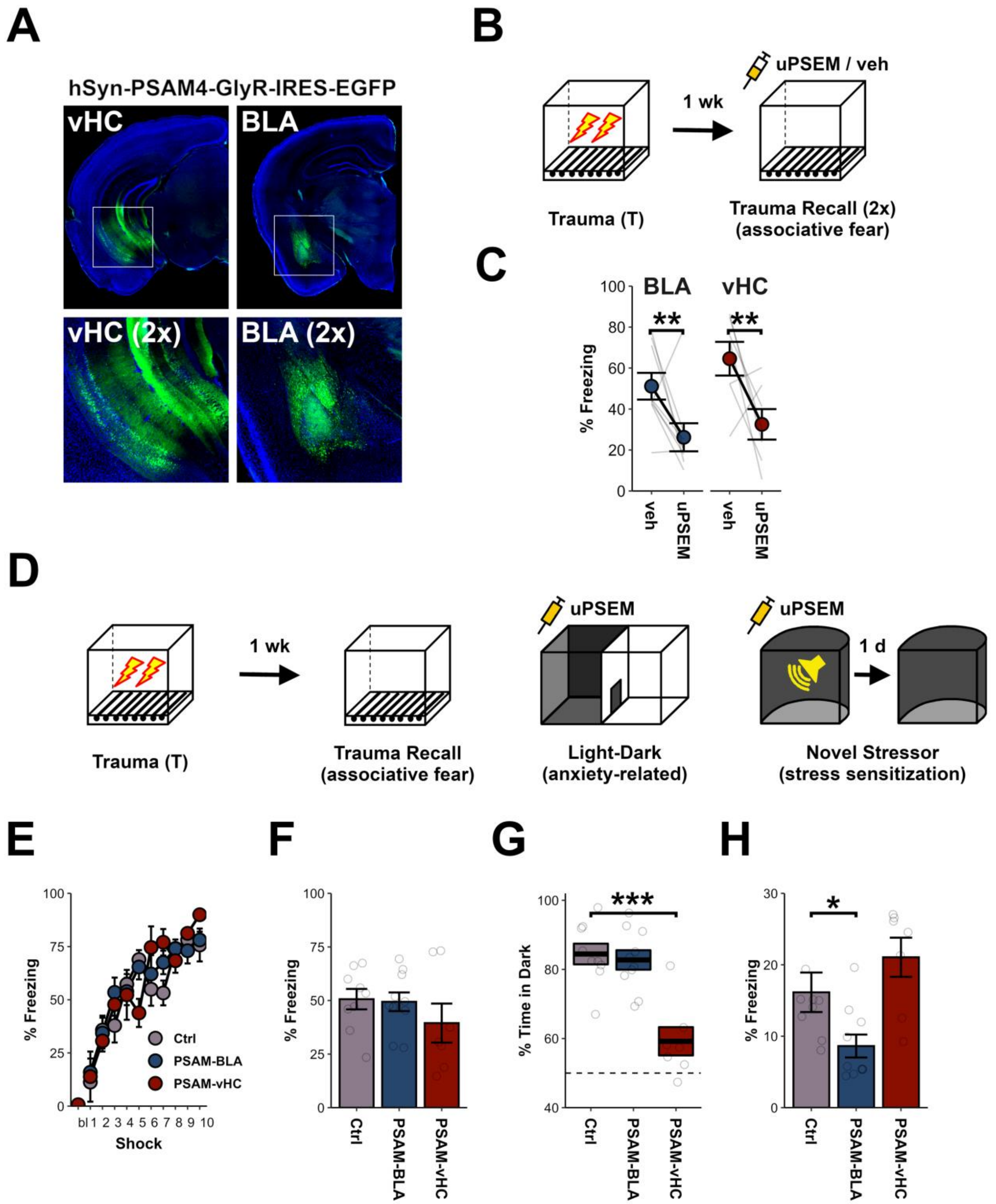


**Figure S8. PSAM inhibition replicates doubly dissociable contributions of BLA and vHC activity to stress sensitization and anxiety-related behavior.**

**A)** A pan-neuronal virus expressing the inhibitory ionotropic receptor PSAM4-GlyR (PSAM) [13] was infused into either the BLA or vHC a month prior to behavioral testing (receptor distribution similar to that in Fig S7).

**B)** A first set of animals received trauma and a week later were tested for trauma recall twice, once after receiving an injection of the PSAM agonist uPSEM-817-tartrate (uPSEM) [13], and once after receiving an injection of vehicle.

**C)** In the trauma recall test, inhibition of either the BLA or vHC via administration of the agonist uPSEM reduced associative freezing (Drug: $F_{1,14}$=12.1, p<0.01; Drug X Region: $F_{1,14}$=0.2, p=0.67). Notably, no prior differences were observed in freezing during the trauma experience for animals with the PSAM receptor in the BLA versus the vHC (Region: $F_{1,14}$=2, p=0.18; Region x Shock: $F_{10,140}$=0.5, p=0.91).

**D)** In a separate set of animals that underwent trauma, we then tested the effects of inhibiting the BLA or vHC during testing of anxiety-related behavior in the light-dark test and administration of the novel stressor. Animals had PSAM-expressing virus infused into the BLA, vHC, or underwent Ctrl surgery in which PBS was infused into the BLA/vHC. The agonist uPSEM was given to all animals prior to the light-dark test and the novel stressor.

**E)** No behavioral differences were observed during the initial trauma, suggesting that expression of the receptor alone had no effect on the acquisition or expression of conditioned freezing (Group: $F_{2,23}$=0.3, p=0.72; Group x Shock: $F_{20,230}$=1.6, p=0.1).

**F)** In a drug-free trauma recall session, there was no difference in freezing between groups ($F_{2,23}$=0.5, p=0.6)

**G)** Inhibition of the vHC produced a dramatic decrease in anxiety-related behavior in the light-dark test, whereas inhibition of the BLA was without effect (vHC: $t_{11.7}$=5, p<0.001. BLA: $t_{16.8}$=0.4, p=0.68).

**H)** Inhibition of the BLA during the novel stressor resulted in reduced freezing the following day (relative to virus-naïve controls), whereas inhibition of the vHC was without effect (BLA: $t_{13}$=2.4, p=0.04. vHC: $t_{13.8}$=1.3, p=0.23)

For B-C, BLA=9 and vHC=7 mice. For D-H, Ctrl=9, PSAM-BLA=10, and PSAM-vHC=7 mice. p<.05 (*), p<0.01 (**), p<0.001 (***). Error bars reflect standard error of the mean.

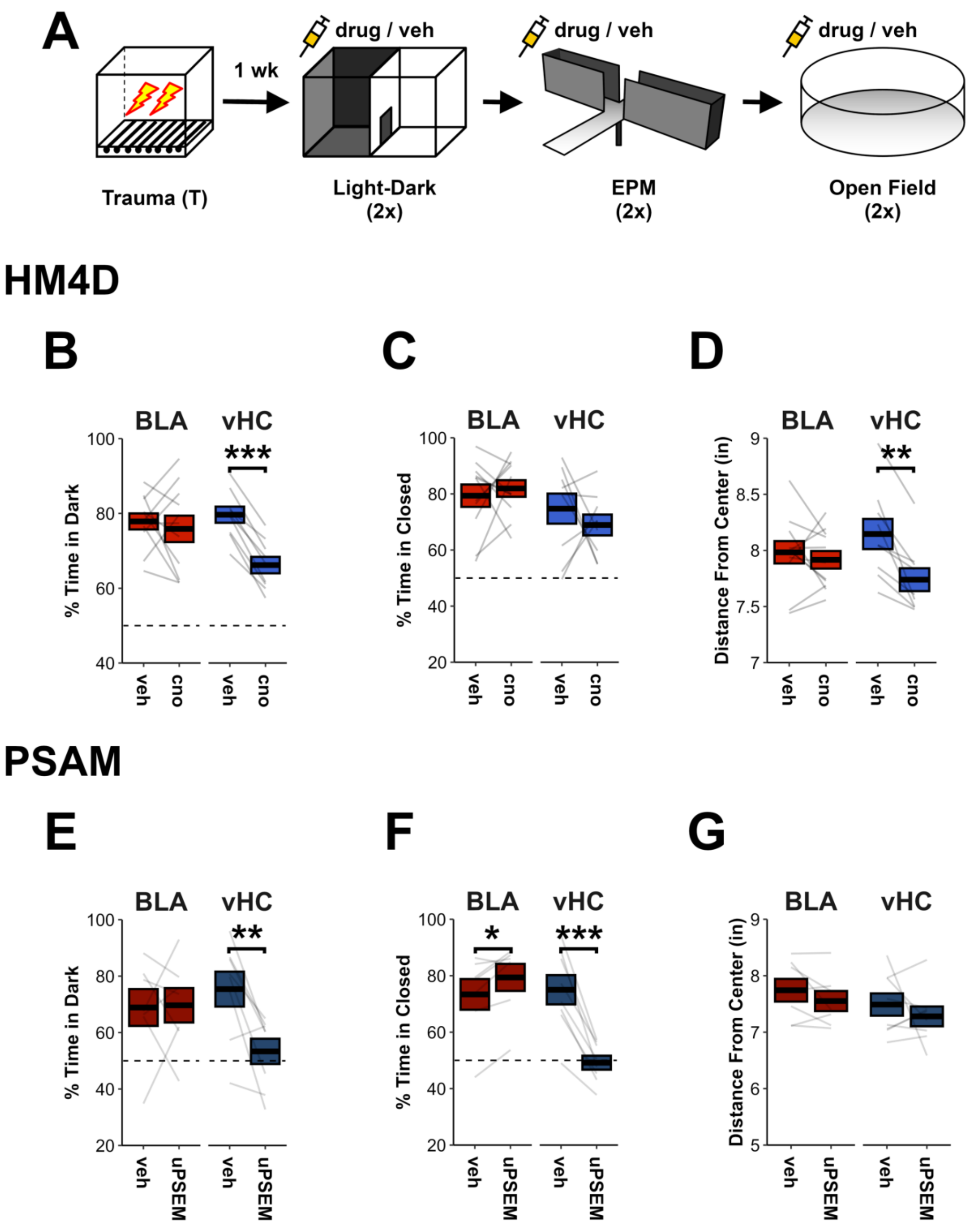


**Figure S9. Activity of the vHC, but not the BLA, is necessary for multiple anxiety-related behaviors.**

**A)** Animals expressing HM4D or PSAM in the BLA/vHC were subjected to trauma and then tested in multiple exploratory anxiety-related tests, both with and without the appropriate chemogenetic agonist (cno for HM4D and uPSEM for PSAM).

**B)** HM4D inhibition of the vHC reduced time in the dark side in the light-dark test, whereas inhibition of the BLA was without effect (vHC: $F_{1,8}$=133.3, p<0.001. BLA: $F_{1,10}$=0.3, p=0.6).

**C)** Niether HM4D inhibition of the vHC nor BLA altered time in the closed arms in the EPM (Drug: $F_{1,18}$=0.1, p=0.71; Region x Drug: $F_{1,18}$=0.93, p=0.34).

**D)** HM4D inhibition of the vHC reduced average distance from the center of the open field, whereas inhibition of the BLA was without effect (vHC: $F_{1,8}$=21, p<0.01. BLA: $F_{1,10}$=0.5, p=0.48).

**E)** PSAM inhibition of the vHC reduced time in the dark side in the light-dark test, whereas inhibition of the BLA was without effect (vHC: $F_{1,7}$=14.9, p<0.01. BLA: $F_{1,6}$=0, p=0.92).

**F)** PSAM inhibition of the vHC reduced time in the closed arms in the EPM, whereas inhibition of the BLA produced a slight increase in time in the closed arms (vHC: $F_{1,7}$=55.6, p=0.001. BLA: $F_{1,6}$=10.2, p=0.02).

**G)** Neither PSAM inhibition of the BLA nor the vHC altered distance from the center in the open field test (Drug: $F_{1,13}$=1.7, p=0.21; Region x Drug: $F_{1,13}$=0, p=0.96).

For HM4D in B-D, BLA=11 and vHC=9 mice. For PSAM in E-G, BLA=7 and vHC=8 mice. p<.05 (*), p<0.01 (**), p<0.001 (***). Error bars reflect standard error of the mean.

### SUPPLEMENTAL METHODS

#### METHODS AND MATERIALS

**Animals***:*

All animals were adult C57BL/6J mice obtained from Jackson Laboratories, aged 2-6 months. A mix of male and female mice were used to demonstrate initial effects of stress on subsequent defensive behavior (Fig 1), as well as the systemic effects of protein synthesis inhibition (Fig 2). Because no sex differences were observed, all other experiments were performed using males. Two animals were excluded from behavioral analyses after they spent >95% time in the light in the light-dark test (Fig 1). All other exclusions were due to inaccurate viral/cannula placement. Animals were housed in a temperature- and humidity-controlled vivarium on a 12/12 light-dark cycle (lights on at 7 a.m.), and all handling and behavioral testing took place during the light phase. All experimental procedures were approved by the Icahn School of Medicine at Mount Sinai's IACUC.

**Behavioral testing:**

For all experiments, animals were singly housed beginning 1 week prior to the start of behavioral testing and were handled by the experimenters for approximately 1 min/day for 5 days during this time. When systemic injections were to be given, animals were additionally briefly habituated to restraint 2-3 times. Animals were habituated to being transported from the vivarium to the laboratory 2-3 times to mitigate transport serving as an associative cue.

*Trauma and trauma recall*: Animals were transported from the vivarium in their cages on a cart to the experimental testing room, which was well lit and had a fan providing ambient sound. Animals were then placed in a brightly lit experimental testing chamber with a grid floor (Med Associates), scented with 5% Simple Green solution. During trauma, after a 5 min period of baseline exploration, animals received 10, 1 sec, 1 mA, scrambled foot-shocks, with an inter-shock interval of 30 sec. Animals were taken out of the testing chamber 30 sec after the last shock and returned to the vivarium in their home cage. For trauma recall sessions, animals were transported to the same experimental testing chamber for an 8 min test session.

*Novel stressor and novel stressor recall*: Animals were transported from the vivarium in P1000 pipet boxes and carried in a dark cardboard box to the experimental testing room, which was dark except for a dim red light. Animals were then placed inside of a dark testing chamber (Med Associates) with a flat plexiglass floor and a curved back wall. The chamber was scented with 1% acetic acid solution.  After a 3 min baseline period, animals were exposed to a single loud auditory stimulus (3 sec, 130 dB white noise, 0 ms rise time) that was delivered by a speaker attached to the wall.  Animals were removed 10 sec later and returned to the vivarium.  For novel stressor recall sessions, animals were transported to the same experimental testing chamber for an 8 min test session.

*Exploratory anxiety-related tests*:  The light-dark test was conducted using two interconnected square compartments with an open top (each compartment measured 7.5 in width x 11.25 in height), separated by a 1.5 in wide passageway that could be closed with an opaque sliding divider.  One chamber was made of all white acrylic, while the walls of the other were covered in matte black wallpaper and had a red acrylic floor.  Overhead lighting provided luminance of 50 lux on the light side.  After a 1 min baseline period in which animals were confined to the dark side, the central divider was raised and the animals could freely explore both sides of the light-dark box.  The open field test was conducted in a circular arena (19 in diameter; 10 in height) made of white acrylic.  A circular field was utilized in order to avoid the need to define arbitrary center/outer areas.  Animals were placed along one wall and allowed to explore for 5 min.  Luminance was approximately 50 lux.  For the elevated plus maze, each arm measured 2 3/8 in wide and 13.75 in long.  The floor of the maze was made of white acrylic, and the enclosed arms had black walls (8 in high).  Luminance of the open arms was 25 lux.  Animals were placed at the end of a closed arm and then allowed to explore freely for 5 min.  For all anxiety-related behavior tests, apparatus were cleaned with 70% ethanol between test sessions and behavior was captured with an overhead webcam.  These sessions were conducted in an otherwise dark room with a fan providing ambient background noise.

*Learning Rate Analysis:* To examine differences in the acquisition of associative fear after trauma (Fig S2), animals experienced the same 10 foot-shock trauma described above, or were placed in the same environment but received no shocks. The next day, mice were placed back in the trauma context for an 8 minute context test.  Beginning the following day, all animals were placed in a novel environment and received one weak shock per day across 7 days (Foot-shock = 2 sec, 0.25 mA.  3 minute baseline.  Taken out 30 sec after shock.  Note this is the same environment across these 7 days).  A low amplitude foot-shock was utilized because it is known to produce lower asymptotic freezing levels [96].  Data is compared qualitatively to predictions from the Rescorla-Wager Model [97].  In addition to looking at the percentage of time spent freezing, we also examined how animals progressed toward asymptotic freezing levels. For each animal, asymptote was defined by their average level of freezing across days 3-7. We then calculated freezing on each day as a percent of asymptotic freezing. Lastly, we assessed shock reactivity evoked by the low intensity shock in the novel environment. For each animal, average baseline motion prior to shock onset, as well as average motion during the shock, was ascertained across the seven days of conditioning. Group differences in shock-induced motion were then assessed.

*Behavior quantification:* For analysis of freezing and motion in conditioning chambers, Med Associates Video Freeze software was used to analyze videos acquired from a near infra-red camera located in the chamber [98]. For measuring distance traveled and time spent in regions of interest in exploratory anxiety-related behavior tests, ezTrack was used [99,100]. With the exception of freezing during the trauma and novel stressor session, all measures reflect the average across the entire session. For freezing during the trauma, time was binned into the 300 sec baseline and then 10 post-shock periods. Each post-shock period was 20 sec in length and began 10 sec after shock offset.

**Surgery:**

For surgery, anesthesia was induced with 5% isoflurane and subsequently maintained at 1-2%. Body temperature was maintained during surgery and recovery with a heating pad below the animal, and ophthalmic ointment was applied to lubricate the eyes. All surgeries followed aseptic surgical technique. For viral surgeries, 100-150 nL was infused into the BLA (AP: -1.4; ML: 3.3; DV: -5) or vHC (AP: -3; ML: 3.2; DV: -4.5) at 2 nL/sec via glass pipettes. For pan-neuronal HM4D experiments, either 100 nL of AAV5-hSyn-HM4Di-mCherry (7 x $10^{12}$ GC/mL; Addgene 50475) or AAV5-hSyn-eGFP (2.26 x $10^{12}$; Addgene 50465) were infused. For projection-specific HM4D experiments, 150 nL containing a cocktail of AAVrg-ef1a-Cre (final concentration=1.1 x $10^{13}$ GC/mL; Addgene 55636) and AAV8-hSyn-eGFP (final concentration=3.8 x $10^{12}$ GC/mL; Addgene 50465) were infused into the projection target structure. Additionally, 150 nL of AAV5-hSyn-DIO-HM4Di-mCherry (2.4 x $10^{13}$ GC/mL; Addgene 44362) or AAV5-hSyn-DIO-mCherry (7.3 x $10^{12}$ GC/mL; Addgene 50459) was infused into the projection origin structure. For PSAM experiments, 100 nL of AAV5-hSyn-PSAM4-GlyR-IRES-EGFP (2.4 x $10^{13}$ GC/mL; Addgene 119742) was infused. Alternatively, an equivalent volume of sterile PBS was infused. 10 min was allowed for diffusion before removing the injector, irrigating the incision with saline, and suturing the incision site. For cannulation surgeries, 26 gauge guide cannula (P1 Technologies; 8IC315GMNSPC) were implanted overlying the BLA (AP: -1.4; ML: 3.2; DV: -3.5) or vHC (AP: -3; ML: 3.2; DV: -3), and affixed to the skull with dental cement and super glue. A skull screw was also implanted during surgery to help secure the head cap (P1 Technologies; 00-96X1/16). After surgery, dummy cannula that extended 1.5 mm below the guide cannula were inserted (P1 Technologies 8IC315DCMNSP). Following surgery, animals were given 20 mg/kg ampicillin and 5 mg/kg carprofen (s.c.) per day for 7 days and body weight and general disposition were monitored. All surgeries followed aseptic surgical technique.

**Anisomycin experiments:**

For experiments in which anisomycin (Sigma A9789) was administered systemically, we utilized a dose of 150 mg/kg (10 mL/kg, s.c.), consistent with prior literature [101,102]. Because numerous waves of protein synthesis have been found to support memory consolidation [68,103,104], we opted to administer anisomycin 3 times, once every 4 hours. In line with prior reports [105,106], this should maintain approximately 90% blockade of protein synthesis for 12 hours. Control animals treated with vehicle were injected/infused at the same times as animals receiving anisomycin (saline for systemic injections; PBS for intracranial injections). For experiments in which anisomycin was administered intracranially, 33 gauge injectors (P1 Technologies; 8IC315IMNSPC) attached via PE-20 tubing (Instech) to a Harvard syringe pump (Harvard Apparatus, #55-2222) were utilized to infuse anisomycin (10 ng/nL) at a rate of 150 nL/min. 300 nL of anisomycin solution was administered per hemisphere in the vHC. 200 nL was administered per hemisphere in the BLA. Control animals were infused with an equivalent volume of 1X PBS. Critically, we used a dose of anisomycin that was unable to affect memory recall (Fig S5), but was nevertheless able to alter translation of the immediate early gene cFos (Fig S5). Moreover, although the dosing regimen used was found to alter memory consolidation when given immediately after a learning event, it had no long-term deleterious impacts on future learning (Fig S6). Following infusions, injectors were left in place for 1 min before removal. Again, anisomycin was infused 3 times, once every four hours. Prior to testing, animals were habituated to handling such that infusions could be done while mice were gently held by the experimenter. Additionally, all animals received a habituation infusion of 1X PBS 2-3 days prior to the trauma day. Anisomycin was first dissolved in a small volume of 0.1 N HCL (90% PBS, 10% 1 N HCL), brought near concentration with the addition of 1X PBS, and the pH was then normalized to 6-7 by the addition of 1 N NaOH.

**HM4D experiments:**

For HM4D experiments, actuation of HM4D was achieved through intraperitoneal administration of 3 mg/kg cno-dihydrochloride (Tocris), 30-40 minutes prior to behavior, at a volume of 10 mL/kg (dissolved in saline). For the study in which the effects of inhibiting the vHC or BLA were assessed on multiple measures of exploratory anxiety-related behavior, each animal underwent each of these tests twice, once with cno and once with vehicle, in a counterbalanced order. Tests occurred in a counterbalanced order, with the constraint that each test (EPM, open field, light-dark) was experienced once before that test was repeated.

**PSAM experiments:**

For PSAM experiments, actuation of PSAM4-GlyR was achieved through intraperitoneal administration of 1 mg/kg uPSEM-817-tartrate (Tocris), 15-20 minutes prior to behavior, at a volume of 10 mL/kg (dissolved in saline). For the study in which the effects of inhibiting the vHC or BLA were assessed on multiple measures of exploratory anxiety-related behavior, each animal underwent each of these tests twice, once with uPSEM and once with vehicle. Tests occurred in a fixed order across two weeks, with open-field on Monday, EPM on Wednesday, and Light-Dark on Friday. However, drug order was counterbalanced, such that half the animals that received uPSEM on the first open field test received saline on the first EPM, and so forth.

**Histology:**

At the end of behavioral testing, animals that underwent surgical manipulation were deeply anesthetized, and their brains were then extracted and placed in paraformaldehyde overnight at 4C. For animals with cannula implants, 100 nl of DAPI (0.5 mg/mL) was infused prior to brain extraction, but after anesthesia, to mark cannula placement. The next day, brains were transferred to 30% sucrose in 1X PBS and left at 4C to sink before being frozen and sectioned at 50 um on a cryostat. Tissue was then mounted on slides and either cover-slipped using mounting media with DAPI (Vector Laboratories, #H-1200-10) for checking viral placement or cover-slipped with non-fluorescent mounting media (Vector Laboratories, #H-1000-10) after a green nucleic acid stain. For green nucleic acid staining, slides were submerged in 50 mM Sytox Green (diluted in 1X PBS from 5 mM. Thermo Fisher #S7020) for 10 min and then washed 3x in 1X PBS. Tissue was then imaged on a Leica DM6 epifluorescent microscope. Viral expression and cannula placement was evaluated using the mouse brain atlas of Franklin and Paxinos [107].

***Ex vivo* electrophysiology:**

Acute coronal brain slices were prepared in slice cutting solution (20 mM NaCl, 3.5 mM KCl, 1.4mM $NaH_2PO_4$, 26 mM $NaHCO_3$, 11 mM D-glucose, 175 mM sucrose, 1.3 mM $MgCl_2$) at a thickness of 350 µm. All recordings for BLA and vHC principal neurons were performed with a MultiClamp 700B amplifier (Molecular Devices) in artificial cerebrospinal fluid solution (119 mM NaCl, 2.5 mM $CaCl_2$, 2.5 mM KCl, 1.25 mM $NaH_2PO_4$, 2 mM $MgSO_4$, 26 mM $NaHCO_3$, and 10 mM D-glucose, equilibrated with 95% $O_2$ and 5% $CO_2$ at pH 7.2 - 7.4) [108]. Fluorescence signal was identified under a Nikon Eclipse FN1 microscope (Nikon Instrument Inc.) using SOLA light engine (Lumencor). Cell-attached recordings were performed in voltage clamp mode. The baseline firing rate was measured 10 minutes prior to bath application of clozapine N-oxide (CNO, 20 µM). The firing rate was then continuously measured over 50 minutes of CNO application. Average firing rate during the last 10 minutes were then compared to the baseline firing rate. Data was analyzed using Clampfit 10.7 software (Molecular Devices).

**Immunohistochemistry and image analysis:**

Immunohistochemistry for cFos was performed on 50 um free-floating tissue sections. In brief, tissue was blocked for one hour in a mixture of 0.3% Triton X-100 and 3% normal goat serum in 1X PBS. Tissue was then incubated in a solution of primary antibody plus blocking solution overnight (Synaptic Systems 226 003, polyclonal rabbit anti cFos, 1:2000), followed by incubation in secondary antibody for ~4 hours (Invitrogen A-11036, goat-anti rabbit Alexa Fluor 568, 1:500). Tissue was washed 3 times, 10 min each, before and after incubation in secondary. All steps were performed at room temperature with gentle shaking. Lastly, tissue was mounted on slides and coverslipped using mounting media with DAPI.

Images were acquired at 10x on a Leica DM6B microscope, using identical exposure settings across animals. For each animal, 4-5 image tiles of the entire amygdala were collected.

c-Fos counts were performed in an automated manner utilizing custom in-house code (https://github.com/ZachPenn/CellCounting). In brief, a median filter was applied to each image to remove granular noise, local fluctuations in background fluorescence were removed with a large gaussian kernel, and a binary threshold was then applied to separate cells from background. A watershed algorithm was implemented to separate adjacent cells and individual cell puncta were then counted. All parameters were applied in an equivalent manner to all images and selected based upon concordance with a set of manually counted images. Regions of interest were drawn and the number of cFos positive cells per unit area was determined.

**Analysis:**

All analyses were performed using RStudio. All data and statistical analysis are available at github.com/ZachPenn/BLAvHC_Dissoc. Group sizes are listed in each figure legend. Briefly, omnibus ANOVA were conducted using the package ezANOVA with type 3 degrees of freedom. The white adjustment was implemented to correct for heterogeneity of variance using heteroscedasticity corrected standard errors (‘hc3’). For repeated measures ANOVA, the Greenhouse-Geisser correction was implemented when the assumption of sphericity was not met. Post-hoc t-tests and planned comparisons did not assume equal variance between groups (Welch test). Post-hoc t-tests were only conducted following main effects/interactions at omnibus significance, but omnibus tests are not reported in the main text for the sake of clarity. Detailed statistics with omnibus results can be found in Supplementary Table 1. Post-hoc tests and planned comparisons were evaluated against a modified criterion calculated using the Dunn-Sidak method in order to keep family-wise type 1 error at 0.05. Criterion p values for significance can be found in Supplementary Table 1. F and t values are rounded to the nearest tenth and hundredth, respectively. Where F values were less than .1, F

### References and Literature Cited


1. Rau, V., DeCola, J.P., and Fanselow, M.S. (2005). Stress-induced enhancement of fear learning: an animal model of posttraumatic stress disorder. Neurosci Biobehav Rev 29, 1207-1223. 10.1016/j.neubiorev.2005.04.010.
2. Rescorla, R.W., and Wagner, A.R. (1972). A theory of Pavlovian conditioning: Variations in the effectiveness of reinforcement and nonreinforcement. In Classical conditioning: II. Current research and theory, B.A. H, and P.W. F, eds. (Appleton-Century-Crofts), pp. 64-99.
3. Ozawa, T., Ycu, E.A., Kumar, A., Yeh, L.F., Ahmed, T., Koivumaa, J., and Johansen, J.P. (2017). A feedback neural circuit for calibrating aversive memory strength. Nat Neurosci 20, 90-97. 10.1038/nn.4439.
4. Franklin, K., and Paxinos, G. (2008). The mouse brain in stereotaxic coordinates, 3 Edition (Elsevier Inc.).
5. Sharma, A.V., Nargang, F.E., and Dickson, C.T. (2012). Neurosilence: profound suppression of neural activity following intracerebral administration of the protein synthesis inhibitor anisomycin. J Neurosci 32, 2377-2387. 10.1523/JNEUROSCI.3543-11.2012.
6. Schafe, G.E., and LeDoux, J.E. (2000). Memory consolidation of auditory pavlovian fear conditioning requires protein synthesis and protein kinase A in the amygdala. J Neurosci 20, RC96.
7. Nader, K., Schafe, G.E., and Le Doux, J.E. (2000). Fear memories require protein synthesis in the amygdala for reconsolidation after retrieval. Nature 406, 722-726. 10.1038/35021052.
8. Frankland, P.W., Ding, H.K., Takahashi, E., Suzuki, A., Kida, S., and Silva, A.J. (2006). Stability of recent and remote contextual fear memory. Learn Mem 13, 451-457. 10.1101/lm.183406.
9. Werka, T., Skår, J., and Ursin, H. (1978). Exploration and avoidance in rats with lesions in amygdala and piriform cortex. J Comp Physiol Psychol 92, 672-681.
10. Maren, S., Aharonov, G., and Fanselow, M.S. (1996). Retrograde abolition of conditional fear after excitotoxic lesions in the basolateral amygdala of rats: absence of a temporal gradient. Behav Neurosci 110, 718-726.
11. Sprengelmeyer, R., Young, A.W., Schroeder, U., Grossenbacher, P.G., Federlein, J., Büttner, T., and Przuntek, H. (1999). Knowing no fear. Proc Biol Sci 266, 2451-2456. 10.1098/rspb.1999.0945.
12. Gale, G.D., Anagnostaras, S.G., Godsil, B.P., Mitchell, S., Nozawa, T., Sage, J.R., Wiltgen, B., and Fanselow, M.S. (2004). Role of the basolateral amygdala in the storage of fear memories across the adult lifetime of rats. J Neurosci 24, 3810-3815. 10.1523/JNEUROSCI.4100-03.2004.
13. Magnus, C.J., Lee, P.H., Bonaventura, J., Zemla, R., Gomez, J.L., Ramirez, M.H., Hu, X., Galvan, A., Basu, J., Michaelides, M., and Sternson, S.M. (2019). Ultrapotent chemogenetics for research and potential clinical applications. Science 364. 10.1126/science.aav5282.
