## Supplemental Table for "Dissociable contributions of the amygdala and ventral hippocampus to stress-induced changes in defensive behavior"

***Notes***

***** = *p<0.05*

****** = *p<0.01*

******* = *p<0.001*

**#** = *nominally p<0.05, but not significant when correcting for multiple comparisons.*

### Figure 1

**Base Behavioral Effects**

**1B**

Trauma Post-Shock Freezing

Repeated Measures Factorial ANOVA (Trauma x Shock x Sex)

| | | |
|---|---|---|
| Trauma: | $F_{1,33}$=167.4 | p<0.001 *** |
| Shock: | $F_{10,330}$=18.7 | p<0.001 *** |
| Sex: | $F_{1,33}$=0.5 | p=0.48 |
| Trauma x Shock: | $F_{10,330}$=16 | p<0.001 *** |
| Trauma x Sex: | $F_{1,33}$=0.6 | p=0.44 |
| Shock x Sex: | $F_{10,330}$=0.8 | p=0.58 |
| Trauma x Shock x Sex: | $F_{10,330}$=0.6 | p=0.7 |

**1C**

Trauma Recall

Factorial ANOVA (Trauma x Sex)

| | | |
|---|---|---|
| Trauma: | $F_{1,52}$=121.6 | p<0.001 *** |
| Sex: | $F_{1,52}$=0 | p=0.97 |
| Trauma x Sex: | $F_{1,52}$=0 | p=0.96 |

**1D**

Light-Dark

Factorial ANOVA (Trauma x Sex)

| | | |
|---|---|---|
| Trauma: | $F_{1,52}$=19.7 | p<0.001 *** |
| Sex: | $F_{1,52}$=1.4 | p=0.91 |
| Trauma x Sex: | $F_{1,52}$=3.3 | p=0.57 |

**1E**

Novel Stressor Baseline

Factorial ANOVA (Trauma x Sex)

| | | |
|---|---|---|
| Trauma: | $F_{1,52}$=2.6 | p=0.11 |
| Sex: | $F_{1,52}$=0.6 | p=0.44 |
| Trauma x Sex: | $F_{1,52}$=0.9 | p=0.35 |

Novel Stressor Test

Factorial ANOVA (Trauma x Sex)

| | | |
|---|---|---|
| Trauma: | $F_{1,52}$=16.1 | p<0.001 *** |
| Sex: | $F_{1,52}$=1.4 | p=0.24 |
| Trauma x Sex: | $F_{1,52}$=0.6 | p0.46 |

**Phenotype Correlations**

**1F**

Trauma Recall x Light-Dark Correlation

| | | |
|---|---|---|
| Pearson correlation: | R=0.21, $t_{38}$=1.3 | p=0.19 |

**1G**

Trauma Recall x Novel Stressor Correlation

| | | |
|---|---|---|
| Pearson correlation: | R=0.44, $t_{38}$=3 | p<0.01 ** |

**1H**

Light-Dark x Novel Stressor Correlation

| | | |
|---|---|---|
| Pearson correlation: | R=0.16, $t_{38}$=1 | p=0.32 |

#### Figure 2

**Systemic Anisomycin**

**2B**

Trauma Post-Shock Freezing (Trauma Groups Only)

Repeated Measures Factorial ANOVA (Group x Shock x Sex)

| | | |
|---|---|---|
| Trauma: | $F_{2,43}=2.8$ | p=0.07 |
| Shock: | $F_{10,430}=92.3$ | p<0.001 *** |
| Sex: | $F_{1,43}=1.7$ | p=0.2 |
| Group x Shock: | $F_{20,430}=0.7$ | p=0.75 |
| Group x Sex: | $F_{2,43}=0.9$ | p=0.42 |
| Shock x Sex: | $F_{10,430}=3.8$ | p<0.001 *** |
| Group x Shock x Sex: | $F_{20,430}=1$ | p=0.45 |

**2C**

Trauma Recall

Factorial ANOVA (Group x Shock x Sex)

| | | |
|---|---|---|
| Group: | $F_{3,58}=33$ | p<0.001 *** |
| Sex: | $F_{1,58}=0$ | p=0.95 |
| Group x Sex: | $F_{3,58}=0.27$ | p=0.85 |

Planned Comparisons (Sidak p-crit=0.017):

| | | |
|---|---|---|
| NT: veh vs. T: veh: | $t_{25}=10.5$ | p<0.001 *** |
| T: veh vs T: ani (0h): | $t_{26.1}=9.5$ | p<0.001 *** |
| T: veh vs T: ani (48h): | $t_{13.3}=0.1$ | p=0.95 |

**2D**

Light-Dark

Factorial ANOVA (Group x Shock x Sex)

| | | |
|---|---|---|
| Group: | $F_{3,58}=4.5$ | p<0.01 ** |
| Sex: | $F_{1,58}=1.5$ | p=0.22 |
| Group x Sex: | $F_{3,58}=0.6$ | p=0.64 |

Planned Comparisons (Sidak p-crit=0.017):

| | | |
|---|---|---|
| NT: veh vs. T: veh: | $t_{33.8}=4.8$ | p<0.001 *** |
| T: veh vs T: ani (0h): | $t_{36.1}=2.8$ | p<0.01 ** |
| T: veh vs T: ani (48h): | $t_{24.7}=0.2$ | p=0.83 |

**2E**

Novel Stressor Test

Factorial ANOVA (Group x Shock x Sex)

| | | |
|---|---|---|
| Group: | $F_{3,58}=5.3$ | p<0.01 ** |
| Sex: | $F_{1,58}=2.6$ | p=0.11 |
| Group x Sex: | $F_{3,58}=0.3$ | p=0.81 |

Planned Comparisons (Sidak p-crit=0.017):

| | | |
|---|---|---|
| NT: veh vs. T: veh: | $t_{29.7}=4.1$ | p<0.001 *** |
| T: veh vs T: ani (0h): | $t_{33.6}=3.4$ | p<0.01 ** |
| T: veh vs T: ani (48h): | $t_{12}=0.7$ | p=0.49 |

**Intracranial Anisomycin**

**2F**

Trauma Post-Shock Freezing (Trauma Groups Only)

Repeated Measures Factorial ANOVA (Group x Shock)

| | | |
|---|---|---|
| Group: | $F_{2,76}$=1.1 | p=0.33 |
| Shock: | $F_{10,760}$=140.5 | p<0.001 *** |
| Group x Shock: | $F_{20,760}$=0.5 | p=0.96 |

**2G**

<u>Trauma Recall</u>

Univariate ANOVA

| | | |
|---|---|---|
| Group: | $F_{3,98}$=49.1 | p<0.001 *** |

Planned Comparisons (Sidak p-crit=0.0085)

| | | |
|---|---|---|
| NT: veh vs. T: veh: | $t_{40.7}$11.5 | p<0.001 *** |
| T: veh vs T: ani-BLA: | $t_{56.9}$=8.2 | p<0.001 *** |
| T: veh vs T: ani-vHC: | $t_{44}$=4.2 | p<0.001 *** |
| T: ani-BLA vs T: ani-vHC: | $t_{30.5}$=2.4 | p=0.02 **#** |
| NT: veh vs T: ani-BLA: | $t_{19.6}$=2.3 | p=0.03 **#** |
| NT: veh vs T: ani-vHC: | $t_{19.6}$=4.1 | p<0.001 *** |

**2H**

<u>Light-Dark</u>

Univariate ANOVA

| | | |
|---|---|---|
| Group: | $F_{3,98}$=11.6 | p<0.001 *** |

Planned Comparisons (Sidak p-crit=0.0085)

| | | |
|---|---|---|
| NT: veh vs. T: veh: | $t_{47.5}$=5.5 | p<0.001 *** |
| T: veh vs T: ani-BLA: | $t_{44.9}$=1.1 | p=0.28 |
| T: veh vs T: ani-vHC: | $t_{28.2}$=3.1 | p<0.01 ** |
| T: ani-BLA vs T: ani-vHC | $t_{29.2}$=0.03 | p=0.03 **#** |
| NT: veh vs T: ani-BLA: | $t_{40}$=4.3 | p<0.001 *** |
| NT: veh vs T: ani-vHC: | $t_{32.2}$=0.8 | p=0.45 |

**2I**

<u>Novel Stressor Test</u>

Univariate ANOVA

| | | |
|---|---|---|
| Group: | $F_{3,98}$=13.1 | p<0.001 *** |

Planned Comparisons (Sidak p-crit=0.0085)

| | | |
|---|---|---|
| NT: veh vs. T: veh: | $t_{47.6}$=5.8 | p<0.001 *** |
| T: veh vs T: ani-BLA: | $t_{56.4}$=4.1 | p<0.001 *** |
| T: veh vs T: ani-vHC: | $t_{46.2}$=1.6 | p=0.12 |
| T: ani-BLA vs T: ani-vHC: | $t_{28.8}$=1.9 | p=0.07 |
| NT: veh vs T: ani-BLA: | $t_{29.1}$=1.9 | p=0.07 |
| NT: veh vs T: ani-vHC: | $t_{22.5}$=3.1 | p<0.01 ** |

### Figure 3

**Slice Electrophysiology**

**3B**

Firing Rate Change

Repeated Measures Factorial ANOVA (All 3 Cell Types x Time)

| | | |
|---|---|---|
| Cell Type: | $F_{2,11}=11.6$ | $p<0.01$ ** |
| Time: | $F_{1,11}=57.3$ | $p<0.001$ *** |
| Cell Type x Time: | $F_{2,11}=11.6$ | $p<0.01$ *** |

Repeated Measures Factorial ANOVA for HM4D+ Cells Only (Region x Time)

| | | |
|---|---|---|
| Region: | $F_{1,10}=0.4$ | $p=0.52$ |
| Time: | $F_{1,10}=137$ | $p<0.001$ *** |
| Region x Time: | $F_{1,10}=0.4$ | $p=0.52$ |

Repeated Measures ANOVA for HM4D- Cells Only

| | | |
|---|---|---|
| Time: | $F_{1,1}=0.2$ | $p=0.76$ |

**BLA/vHC Inhibition on Associative Fear Recall**

**3D**

Trauma Post-Shock Freezing

Repeated Measures Factorial ANOVA (Region x Shock)

| | | |
|---|---|---|
| Region: | $F_{1,28}=0.3$ | $p=0.59$ |
| Shock: | $F_{10,280}=71.8$ | $p<0.001$ |
| Region x Shock: | $F_{10,280}=0.7$ | $p=0.66$ |

**3E**

Inhibition on Trauma Recall Test

Repeated Measures Factorial ANOVA (Region x Drug x Time)

| | | |
|---|---|---|
| Region: | $F_{1,26}=0.3$ | $p=0.57$ |
| Drug: | $F_{1,26}=0.7$ | $p=0.41$ |
| Time: | $F_{1,26}=31.2$ | $p<0.001$ *** |
| Region x Drug: | $F_{1,26}=0$ | $p=0.99$ |
| Region x Time: | $F_{1,26}=5$ | $p=0.03$ |
| Drug x Time | $F_{1,26}=14.7$ | $p<0.001$ *** |
| Region x Drug x Time: | $F_{1,26}=0.4$ | $p=0.51$ |

Repeated Measures Factorial ANOVA for CNO Groups Only (Region x Time)

| | | |
|---|---|---|
| Region: | $F_{1,14}=0.2$ | $p=0.67$ |
| Time: | $F_{1,14}=33.5$ | $p<0.001$ *** |
| Region x Time: | $F_{1,14}=3.2$ | $p=0.1$ |

Repeated Measures Factorial ANOVA for VEH Groups Only (Region x Time)

| | | |
|---|---|---|
| Region: | $F_{1,12}=0.1$ | $p=0.7$ |
| Time: | $F_{1,12}=2.8$ | $p=0.12$ |
| Region x Time: | $F_{1,12}=2.2$ | $p=0.16$ |

**BLA/vHC Inhibition on Light-Dark and Novel Stressor**

**3G**

Trauma Post-Shock Freezing (Drug Free)

Repeated Measures Factorial ANOVA (Group x Shock)

Group: $F_{2,65}=0.2$ p=0.82
Shock: $F_{10,650}=196.3$ p<0.001 ***
Group x Shock: $F_{20,650}=0.6$ p=0.84

**3H**

Trauma Recall (Drug Free)

Univariate ANOVA

Group: $F_{2,65}=0.1$ p=0.88

**3I**

Light-Dark (CNO for All)

Univariate ANOVA

Group: $F_{2,65}=6.6$ p<0.01 **

Planned Comparisons (Sidak p-crit=0.025):

eGFP: BLA/vHC vs HM4D: BLA: $t_{45.8}=0.4$ p=0.71
eGFP: BLA/vHC vs HM4D: vHC: $t_{29.2}=3.3$ p<0.01 **

**3J**

Novel Stressor Test (CNO for All During Novel Stressor, but not at Test)

Univariate ANOVA

Group: $F_{2,64}=4.3$ p=0.02 *

Planned Comparisons (Sidak p-crit=0.025):

eGFP: BLA/vHC vs HM4D: BLA: $t_{41.7}=2.8$ p<0.01 **
eGFP: BLA/vHC vs HM4D: vHC: $t_{40.4}=0.7$ p=0.52

**Comparisons of Animals with eGFP in BLA vs vHC for 3I-J**

Trauma Post-Shock Freezing (Drug Free)

Repeated Measures Factorial ANOVA (Region x Shock)

Region: $F_{1,23}=0.2$ p=0.66
Shock: $F_{10,230}=67.9$ p<0.001 ***
Region x Shock: $F_{10,230}=0.7$ p=0.66

Trauma Recall (Drug Free)

Univariate ANOVA

Region: $F_{1,23}=0.9$ p=0.35

Light-Dark (CNO for All)

Univariate ANOVA

Region: $F_{1,23}=1.9$ p=0.18

Novel Stressor Test (CNO for All During Novel Stressor, but not at Test)

Univariate ANOVA

Region: $F_{1,23}=0.2$ p=0.64

### Figure 4

**Projection-specific Inhibition of BLA→vHC and vHC→BLA on Light-Dark and Novel Stressor**

**4C**

Trauma Post-Shock Freezing

Repeated Measures Factorial ANOVA (Group x Shock)

| | | |
|---|---|---|
| Group: | $F_{2,30}=0.1$ | $p=0.91$ |
| Shock: | $F_{10,300}=58.3$ | $p<0.001$ *** |
| Group x Shock: | $F_{20,300}=0.7$ | $p=0.74$ |

**4D**

Trauma Recall (Drug Free)

Univariate ANOVA

| | | |
|---|---|---|
| Group: | $F_{2,30=}1.9$ | $p=0.17$ |

**4E**

Light-Dark (CNO for All)

Univariate ANOVA

| | | |
|---|---|---|
| Group: | $F_{2,30}=1.1$ | $p=0.35$ |

**4F**

Novel Stressor Test (CNO for All During Novel Stressor, but not at Test)

Univariate ANOVA

| | | |
|---|---|---|
| Group: | $F_{2,30}=2.2$ | $p=0.13$ |

**Comparisons of Animals with mCherry in BLA→vHC vs vHC→BLA for Figure 4**

Trauma Post-Shock Freezing (Drug Free)

Repeated Measures Factorial ANOVA (Projection Origin x Shock)

| | | |
|---|---|---|
| Origin: | $F_{1,9}=3.1$ | $p=0.1$ |
| Shock: | $F_{10,90}=19$ | $p<0.001$ *** |
| Origin x Shock: | | |

Trauma Recall (Drug Free)

Univariate ANOVA

| | | |
|---|---|---|
| Origin: | $F_{1,9}=1.4$ | $p=0.26$ |

Light-Dark (CNO for All)

Univariate ANOVA

| | | |
|---|---|---|
| Origin: | $F_{1,9}=0.7$ | $p=0.42$ |

Novel Stressor Test (CNO for All During Novel Stressor, but not at Test)

Univariate ANOVA

| | | |
|---|---|---|
| Origin: | $F_{1,9}=2.5$ | $p=0.15$ |

### Supplementary Figure 1

**Effects of Trauma Amplitude and Sex on Base Behavioral Effects**

**S1B**

Trauma Post-Shock Freezing

Repeated Measures Factorial ANOVA (Group x Shock x Sex)

| | | |
|---|---|---|
| Group: | $F_{2,60}=136.9$ | $p<0.001$ *** |
| Shock: | $F_{10,600}=32$ | $p<0.001$ *** |
| Sex: | $F_{1,60}=0$ | $p=0.91$ |
| Group x Shock: | $F_{20,600}=14$ | $p<0.001$ *** |
| Group x Sex: | $F_{2,60}=1$ | $p=0.39$ |
| Shock x Sex: | $F_{10,600}=0.73$ | $p=0.65$ |
| Group x Shock x Sex: | $F_{20,600}=1.3$ | $p=0.23$ |

Trauma Post-Shock Freezing (Trauma Groups Only)

Repeated Measures Factorial ANOVA (Group x Shock x Sex)

| | | |
|---|---|---|
| Group: | $F_{1,37}=31.9$ | $p<0.001$ *** |
| Shock: | $F_{10,370}=25.8$ | $p<0.001$ *** |
| Sex: | $F_{1,37}=0$ | $p=0.88$ |
| Group x Shock: | $F_{10,370}=2.4$ | $p<0.01$ ** |
| Group x Sex: | $F_{1,37}=0.9$ | $p=0.34$ |
| Shock x Sex: | $F_{10,370}=0.5$ | $p=0.86$ |
| Group x Shock x Sex: | $F_{10,370}=1.3$ | $p=0.25$ |

**S1C**

Trauma Recall

Factorial ANOVA (Group x Sex)

| | | |
|---|---|---|
| Group: | $F_{2,60}=75.3$ | $p<0.001$ *** |
| Sex: | $F_{1,60}=0.3$ | $p=0.56$ |
| Trauma x Sex: | $F_{2,60}=0.3$ | $p=0.73$ |

Planned Comparisons (Sidak p-crit= 0.025):

| | | |
|---|---|---|
| NT vs T: Low: | $t_{9.3}=6.3$ | $p<0.001$ |
| T:Low vs T: High: | $t_{21}=2.3$ | $p=0.03$ **#** |

**S1D**

Light-Dark

Factorial ANOVA (Group x Sex)

| | | |
|---|---|---|
| Group: | $F_{2,60}=9.9$ | $p<0.001$ *** |
| Sex: | $F_{1,60}=0$ | $p=0.83$ |
| Trauma x Sex: | $F_{2,60}=0.2$ | $p=0.8$ |

Planned Comparisons (Sidak p-crit= 0.025):

| | | |
|---|---|---|
| NT vs T: Low: | $t_{14.1}=1.3$ | $p=0.2$ |
| T:Low vs T: High: | $t_{14.5}=1.8$ | $p=0.1$ |

**S1E**

Novel Stressor Baseline

Factorial ANOVA (Group x Sex)

| | | |
|---|---|---|
| Group: | $F_{2,60}=0.2$ | $p=0.19$ |
| Sex: | $F_{1,60}=0.5$ | $p=0.48$ |
| Trauma x Sex: | $F_{2,60}=0.5$ | $p=0.6$ |

Novel Stressor Test

Factorial ANOVA (Group x Sex)

| | | |
|---|---|---|
| Group: | $F_{2,60}=8.7$ | $p<0.001$ *** |
| Sex: | $F_{1,60}=0.4$ | $p=0.52$ |
| Trauma x Sex: | $F_{1,60}=0.6$ | $p=0.56$ |

Planned Comparisons (Sidak p-crit= 0.025):

| | | |
|---|---|---|
| NT vs T: Low: | $t_{14.7}=2.1$ | $p=0.06$ |
| T:Low vs T: High: | $t_{33.1}=2.1$ | $p=0.5$ |

### Supplementary Figure 2

#### Trauma impacts learning consistent with changes in stimulus sensitivity

**S2C**

Pre-Shock Freezing Across 7 Days of Weak Shock

Repeated Measures Factorial ANOVA (Trauma x Day)

| | | |
|---|---|---|
| Trauma: | $F_{1,22}=10.2$ | $p<0.01$ ** |
| Day: | $F_{6,132}=20$ | $p<0.001$ *** |
| Trauma x Day: | $F_{6,132}=1.4$ | $p=0.23$ |

**S2D**

Normalized (to Asymptote) Pre-Shock Freezing Across 7 Days of Weak Shock

Repeated Measures Factorial ANOVA (Trauma x Day)

| | | |
|---|---|---|
| Trauma: | $F_{1,22}=0.8$ | $p=0.39$ |
| Day: | $F_{6,132}=16.6$ | $p<0.001$ *** |
| Trauma x Day: | $F_{6,132}=0.4$ | $p=0.77$ |

**S2E**

Average Shock Reactivity Across 7 Days of Weak Shock

Repeated Measures Factorial ANOVA (Trauma x Shock)

| | | |
|---|---|---|
| Trauma: | $F_{1,22}=0.3$ | $p=0.62$ |
| Time: | $F_{1,22}=102$ | $p<0.001$ *** |
| Trauma x Time: | $F_{1,22}=1.2$ | $p=0.28$ |

### Supplementary Figure 4

**No differences in animals receiving cannula infusions of vehicle in BLA versus vHC**

**S4B**

Trauma Post-shock Freezing

Repeated Measures Factorial ANOVA (Trauma x Region x Shock)

| | | |
|---|---|---|
| Trauma: | $F_{1,59}=1.9$ | $p<0.001$ *** |
| Region: | $F_{1,59}=0$ | $p=0.85$ |
| Shock: | $F_{10,590}=4$ | $p<0.001$ *** |
| Trauma x Shock: | $F_{10,590}=3.8$ | $p<0.001$ *** |
| Trauma x Region: | $F_{1,59}=0$ | $p=0.99$ |
| Shock x Region: | $F_{10,590}=0.6$ | $p=0.75$ |
| Trauma x Region x Shock: | $F_{10,590}=0.8$ | $p=0.56$ |

**S4C**

Trauma Recall

Factorial ANOVA (Trauma x Region)

| | | |
|---|---|---|
| Trauma: | $F_{1,59}=126.6$ | $p<0.001$ *** |
| Region: | $F_{1,59}=0.9$ | $p=0.35$ |
| Trauma x Region: | $F_{1,59}=0.6$ | $p=0.46$ |

**S4D**

Light-Dark

Factorial ANOVA (Trauma x Region)

| | | |
|---|---|---|
| Trauma: | $F_{1,59}=2.7$ | $p<0.001$ *** |
| Region: | $F_{1,59}=0.7$ | $p=0.4$ |
| Trauma x Region: | $F_{1,59}=0.1$ | $p=0.75$ |

**S4E**

Novel Stressor Test

Factorial ANOVA (Trauma x Region)

| | | |
|---|---|---|
| Trauma: | $F_{1,59}=3.1$ | $p<0.001$ *** |
| Region: | $F_{1,59}=0.4$ | $p=0.55$ |
| Trauma x Region: | $F_{1,59}=0$ | $p=0.98$ |

### Supplementary Figure 5

**Effects of anisomycin in BLA on fear memory acquisition**

**S5B**

Baseline Freezing

Univariate ANOVA

| | | |
|---|---|---|
| Drug: | $F_{1,7}=0.3$ | $p=0.62$ |

Post-Shock Freezing

Repeated Measures Factorial ANOVA (Drug x Shock)

| | | |
|---|---|---|
| Drug: | $F_{1,7}=0.3$ | $p=0.58$ |
| Shock: | $F_{2,14}=7.6$ | $p<0.01$ ** |
| Drug x Shock: | $F_{2,14}=0.3$ | $p=0.72$ |

Recall Test Freezing

Univariate ANOVA

| | | |
|---|---|---|
| Drug: | $F_{1,7}=0.3$ | $p=0.62$ |

**Effects of anisomycin in BLA on fear memory recall**

**S5D**

Anisomycin on Fear Recall

Repeated Measures Factorial ANOVA (Drug x Test)

| | | |
|---|---|---|
| Drug: | $F_{1,7}=0.5$ | $p=0.5$ |
| Test: | $F_{1,7}=0.4$ | $p=0.03$ * |
| Drug x Test: | $F_{1,7}=13.6$ | $p<0.01$ |

**Effect of anisomycin in BLA on cFos induction by shock**

**S5F**

Anisomcyin on cFOS

Univariate ANOVA

| | | |
|---|---|---|
| Drug: | $F_{1,5}=7.3$ | $p=0.04$ * |

#### Supplementary Figure 6

**Lasting effects of anisomycin in BLA on fear memory acquisition**

**S6B**

Trauma 2 Baseline Freezing:

Univariate ANOVA

| | | |
|---|---|---|
| Group: | $F_{1,10}=0.2$ | $p=0.64$ |

Trauma 2 Post-shock Freezing

Repeated Measures Factorial ANOVA (Group x Shock)

| | | |
|---|---|---|
| Group: | $F_{1,10}=4.3$ | $p=0.07$ |
| Shock: | $F_{9,90}=5$ | $p<0.01$ ** |
| Group x Shock: | $F_{9,90}=0.7$ | $p=0.73$ |

**S6C**

Trauma 2 Recall:

Univariate ANOVA

| | | |
|---|---|---|
| Group: | $F_{1,10}=0.1$ | $p=0.83$ |

#### Supplementary Figure 8

**Lasting effects of anisomycin in BLA on fear memory acquisition**

**S8C**

Trauma Post-Shock Freezing (Drug Free)
Repeated Measures Factorial ANOVA (Group x Shock)

| | | |
|---|---|---|
| Group: | $F_{1,14}=2$ | $p=0.18$ |
| Shock: | $F_{10,140}=34.6$ | $p<0.001$ *** |
| Group x Shock: | $F_{10,140}=0.5$ | $p=0.91$ |

Inhibition on Trauma Recall Test
Repeated Measures Factorial ANOVA (Region x Drug)

| | | |
|---|---|---|
| Region: | $F_{1,14}=2.6$ | $p=0.13$ |
| Drug: | $F_{1,14}=12.1$ | $p<0.01$ ** |
| Region x Drug: | $F_{1,14}=0.2$ | $p=0.67$ |

**BLA/vHC Inhibition on Light-Dark and Novel Stressor**

**S8E**

Trauma Post-Shock Freezing (Drug Free)
Repeated Measures Factorial ANOVA (Group x Shock)

| | | |
|---|---|---|
| Group: | $F_{2,23}=0.3$ | $p=0.72$ |
| Shock: | $F_{10,230}=61.5$ | $p<0.001$ *** |
| Group x Shock: | $F_{20,230}=1.6$ | $p=0.1$ |

**S8F**

Trauma Recall (Drug Free)
Univariate ANOVA

| | | |
|---|---|---|
| Group: | $F_{2,23}=0.5$ | $p=0.6$ |

**S8G**

Light-Dark (CNO for All)
Univariate ANOVA

| | | |
|---|---|---|
| Group: | $F_{2,23}=12.3$ | $p<0.001$ *** |

Planned Comparisons (Sidak p-crit=0.025):

| | | |
|---|---|---|
| Ctrl vs PSAM-BLA: | $t_{16.8}=0.4$ | $p=0.68$ |
| Ctrl vs PSAM-vHC: | $t_{11.7}=5$ | $p<0.001$ *** |

**S8H**

Novel Stressor Test (CNO for All During Novel Stressor, but not at Test)
Univariate ANOVA

| | | |
|---|---|---|
| Group: | $F_{2,23}=7.6$ | $p<0.01$ ** |

Planned Comparisons (Sidak p-crit=0.025):

| | | |
|---|---|---|
| Ctrl vs PSAM-BLA: | $t_{13}=2.4$ | $p=0.035$ **#** |
| Ctrl vs PSAM-vHC: | $t_{13.8}=1.3$ | $p=0.23$ |

**Comparisons of Ctrl Animals with PBS in BLA vs vHC for S8F-H**

Trauma Recall (Drug Free)
Univariate ANOVA

| | | |
|---|---|---|
| Region: | $F_{1,7}=1.6$ | $p=0.25$ |

Light-Dark (CNO for All)

Univariate ANOVA

| | | |
|---|---|---|
| Region: | $F_{1,7}=1.5$ | $p=0.26$ |

Novel Stressor Test (CNO for All During Novel Stressor, but not at Test)

Univariate ANOVA

| | | |
|---|---|---|
| Region: | $F_{1,7}=2.5$ | $p=0.15$ |

#### Supplementary Figure 9

**Effects of inhibiting BLA/vHC on anxiety-related behavior using HM4D**

**S9B**

Light-Dark

Repeated Measures Factorial ANOVA (Region x Drug)

| | | |
|---|---|---|
| Region: | $F_{1,18}=1.6$ | $p=0.23$ |
| Drug: | $F_{1,18}=13.5$ | $p<0.01$ ** |
| Region x Drug: | $F_{1,18}=7.4$ | $p=0.01$ * |

Light-Dark BLA

Repeated Measures ANOVA:

| | | |
|---|---|---|
| Drug: | $F_{1,10}=0.3$ | $p=0.6$ |

Light-Dark vHC

Repeated Measures ANOVA:

| | | |
|---|---|---|
| Drug: | $F_{1,8}=133.3$ | $p<0.001$ *** |

**S9C**

EPM

Repeated Measures Factorial ANOVA (Region x Drug)

| | | |
|---|---|---|
| Region: | $F_{1,18}=5.8$ | $p=0.03$ * |
| Drug: | $F_{1,18}=0.1$ | $p=0.71$ |
| Region x Drug: | $F_{1,18}=0.93$ | $p=0.34$ |

**S9D**

Open Field

Repeated Measures Factorial ANOVA (Region x Drug)

| | | |
|---|---|---|
| Region: | $F_{1,18}=0$ | $p=0.96$ |
| Drug: | $F_{1,18}=1.3$ | $p<0.01$ ** |
| Region x Drug: | $F_{1,18}=6.9$ | $p=0.02$ * |

Open Field BLA

Repeated Measures ANOVA:

| | | |
|---|---|---|
| Drug: | $F_{1,10}=0.5$ | $p=0.48$ |

Open Field vHC

Repeated Measures ANOVA:

| | | |
|---|---|---|
| Drug: | $F_{1,8}=21$ | $p<0.01$ ** |

**Effects of inhibiting BLA/vHC on anxiety-related behavior using PSAM**

**9E**

Light-Dark

Repeated Measures Factorial ANOVA (Region x Drug)

| | | |
|---|---|---|
| Region: | $F_{1,13}=0.5$ | $p=0.48$ |
| Drug: | $F_{1,13}=5.1$ | $p=0.04$ * |
| Region x Drug: | $F_{1,13}=5.8$ | $p=0.03$ * |

Light-Dark BLA

Repeated Measures ANOVA:

| | | |
|---|---|---|
| Drug: | $F_{1,6}=0$ | $p=0.92$ |

Light-Dark vHC

Repeated Measures ANOVA:

| | | |
|---|---|---|
| Drug: | $F_{1,7}=14.9$ | $p<0.01$ ** |

**S9F**

EPM

Repeated Measures Factorial ANOVA (Region x Drug)

| | | |
|---|---|---|
| Region: | $F_{1,13}=5.4$ | $p=0.04$ * |
| Drug: | $F_{1,13}=23.2$ | $p<0.001$ *** |
| Region x Drug: | $F_{1,13}=59.9$ | $p<0.001$ *** |

EPM BLA

Repeated Measures ANOVA:

| | | |
|---|---|---|
| Drug: | $F_{1,6}=10.2$ | $p=0.02$ * |

EPM vHC

Repeated Measures ANOVA:

| | | |
|---|---|---|
| Drug: | $F_{1,7}=55.6$ | $p=0.001$ *** |

**S9G**

Open Field

Repeated Measures Factorial ANOVA (Region x Drug)

| | | |
|---|---|---|
| Region: | $F_{1,13}=1.4$ | $p=0.25$ |
| Drug: | $F_{1,13}=1.7$ | $p=0.21$ |
| Region x Drug: | $F_{1,13}=0$ | $p=0.96$ |
