## Supplementary material for "Dissociable contributions of the amygdala and ventral hippocampus to stress-induced changes in defensive behavior": Key Resources

### KEY RESOURCES FOR:

**KEY RESOURCE TABLE**

| REAGENT or RESOURCE | SOURCE | IDENTIFIER |
|---|---|---|
| **Bacterial and virus strains** | | |
| AAV5-hSyn-hM4Di-mCherry | | Addgene: 50475 |
| AAV5-hSyn-eGFP | | Addgene: 50465 |
| AAV5-hSyn-DIO-HM4Di-mCherry | | Addgene: 44362 |
| AAV5-hSyn-DIO-mCherry | | Addgene: 50459 |
| AAVrg-ef1a-Cre | | Addgene: 55636 |
| AAV8-hSyn-eGFP | | Addgene: 50465 |
| AAV5-hSyn-PSAM4-GlyR-IRES-EGFP | Magnus et al, 2019[76] | Addgene: 119742 |
| **Antibodies** | | |
| Polyclonal rabbit anti cFos | | Synaptic Systems: 226 003 |
| Goat anti rabbit Alexa Fluor 568 | | Invitrogen: A-11036 |
| **Chemicals, peptides, and recombinant proteins** | | |
| Anisomycin from *Streptomyces griseolus* | Millipore Sigma | Millipore Sigma: A9789 |
| CNO-dihydrochloride | Tocris | Tocris: 6329 |
| uPSEM 817 tartrate | Tocris | Tocris: 6866 |
| **Experimental models: Organisms/strains** | | |
| Mouse: C57BL/6J | The Jackson Laboratory | JAX: 000664 |
| **Software and algorithms** | | |
| Med Associates Video Freeze | Med Associates[98] | Med Associates: SOF-843 |
| ezTrack | Pennington et al, 2019[100] | www.github.com/denisecailab/eztrack |
| Automated cell counting | | www.github.com/zachpenn/cellcounting |
